# Adolescent binge drinking leads to long-lasting changes in cortical microcircuits in mice

**DOI:** 10.1101/2022.08.02.502367

**Authors:** Avery R. Sicher, William D. Starnes, Keith R. Griffith, Nigel C. Dao, Grace C. Smith, Dakota F. Brockway, Nicole A. Crowley

**Affiliations:** Neuroscience Graduate Program, The Huck Institutes of the Life Sciences, University Park PA 16802; Department of Biology, Penn State University, University Park PA 16802; Department of Biomedical Engineering, Penn State University, University Park PA 16802; Center for Neural Engineering, Penn State University, University Park PA 16802

## Abstract

Adolescent drug consumption has increased risks to the individual compared to consumption in adulthood, due to the likelihood of long-term and permanent behavioral and neurological adaptations. However, little is known about how adolescent alcohol consumption influences the maturation and trajectory of cortical circuit development. Here, we explore the consequences of adolescent binge drinking on somatostatin (SST) neuronal function in superficial layers of the prelimbic (PL) cortex in male and female SST-Ai9 mice. We find that adolescent drinking-in-the-dark (DID) produces sex-dependent increases in intrinsic excitability of SST neurons, with no change in overall SST cell number, persisting well into adulthood. While we did not find evidence of altered GABA release from SST neurons onto other neurons within the circuit, we found a complementary reduction in layer II/III pyramidal neuron excitability immediately after binge drinking; however, this hypoexcitability rebounded towards increased pyramidal neuron activity in adulthood in females, suggesting long-term homeostatic adaptations in this circuit. Together, this suggests that binge drinking during key developmental timepoints leads to permanent changes in PL microcircuitry function, which may have broad behavioral implications.

## 1. INTRODUCTION

### 1.1 Adolescence and alcohol misuse

Adolescence is broadly defined as the transitional period between childhood and adulthood, marked by a variety of maturational changes throughout the body (Spear 2000). In the brain, the prefrontal cortex (PFC) is one of the last regions to finalize development, with a variety of morphological, functional, and neuromodulatory changes occurring throughout this period (Sicher et al., 2022). This critical and time-locked development may make the PFC a particularly vulnerable brain region to insults including alcohol use, as any events which interfere with typical PFC development may produce permanent and lifelong consequences. Unfortunately coinciding with this developmental window, adolescents are at a high risk of initiating drug and alcohol use. Alcohol consumption in adolescents often occurs as an episode of binge drinking, defined by the National Institute on Alcohol Abuse and Alcoholism (NIAAA) as a pattern of drinking which raises the blood ethanol concentration (BEC) above 80 mg / dL (NIAAA, 2021). In adult women and men, this typically requires 4 or 5 alcoholic drinks respectively within a 2 hr period; however, adolescents may reach BECs exceeding 80 mg / dL after consuming as few as 3 drinks due to their smaller body size (Donovan, 2009). The NIAAA estimates that around 90% of alcohol consumed by people under the age of 21 is consumed in an episode of binge drinking (NIAAA, 2021)– setting the stage for long-term consequences driven by the adolescent onset of alcohol consumption.

The risks and consequences of adolescent binge drinking are both acute and long-term. While acute risks include car accidents or alcohol poisoning, long-term consequences can include impaired development of the PFC. Studies in both rodent models and human adolescents have shown that alcohol consumption interferes with typical maturation of the prelimbic (PL) region of the PFC, potentially inducing permanent changes in PFC structure and corresponding behavioral aberrations (for review see Sicher et al., 2022). The effects of alcohol on the PL cortex seem to be age dependent, as alcohol consumption during early adolescence, but not early adulthood, increased the excitability of deep-layer cortical pyramidal neurons (Galaj et al., 2020). Similarly, alcohol consumption in adolescence altered medial PFC plasticity, causing persistent changes in fear extinction, whereas adult mice which consumed alcohol showed no behavioral changes (Lawson et al., 2022). This suggests critical windows for the interaction between development, drug exposure, and PFC circuits.

### 1.2 Prelimbic somatostatin neurons and drug use

Somatostatin (SST) is a neuropeptide which represents a promising therapeutic target for substance use disorders and other neuropsychiatric conditions (Brockway and Crowley, 2020). SST is co-expressed in a major population of gamma-aminobutyric acidergic (GABAergic) neurons, with SST neurons representing about 20% of GABAergic neurons in the mouse frontal cortex (Xu et al., 2010). Little work thus far has characterized changes in SST neurons throughout development. During early adolescence, there is an increase in the number of SST neurons in the PL region of the PFC, followed by pruning during the later stages of adolescence in female rodents (Du et al., 2018). Membrane properties of medial PFC (infralimbic and anterior cingulate cortex) SST neurons stabilize early in life, around the third postnatal week (Koppensteiner et al., 2019; Pan et al., 2016), though typical changes in the electrophysiological properties of prelimbic SST neurons during adolescence have not been studied.

Recent work has sought to understand the effects of alcohol on SST neurons. Our lab and others have demonstrated that SST neurons in the PL cortex are vulnerable to several paradigms of alcohol exposure in adult mice (Dao et al., 2021; Joffe et al., 2020). SST has also been identified as an important target of adult alcohol consumption, with SST gene expression correlating with alcohol-induced changes in local circuitry and functional connectivity in humans (Ochi et al., 2022). Together, preclinical and clinical evidence indicate SST neurons are vulnerable to alcohol consumption in adults and may represent a potential therapeutic target for excessive alcohol consumption. However, limited work has explored the effects of alcohol exposure in adolescence on SST neurons. In this study, we investigated the effects of adolescent binge drinking on PL cortical SST neurons, and how changes in PL SST neuronal properties may lead to compensatory changes on downstream pyramidal neurons within the PL cortex.

## 2. MATERIALS AND METHODS

### 2.1 Animals

All experiments were approved by the Pennsylvania State University Institutional Animal Care and Use Committee. Male and female SST-IRES-Cre (stock #013044, The Jackson Laboratory), Ai9 (stock #007909, The Jackson Laboratory), and Ai32 (stock #024109, The Jackson Laboratory) mice were bred in-house. At post-natal day (PND) 21, SST-IRES-Cre:Ai9 and SST-IRES-Cre:Ai32 mice were weaned into single housing and moved into a temperature- and humidity-controlled reverse light cycle room (lights off at 7:00 am) to acclimate for one week before experiments. Littermates were randomly assigned to either alcohol or control conditions as described below. Mice had *ad libitum* access to food and water, except during alcohol exposure.

### 2.2 Adolescent Drinking in the Dark (DID)

DID was conducted as previously published (Dao et al., 2021; Rhodes et al., 2005; Suresh Nair et al., 2022), at timepoints adapted for adolescence (Holstein et al., 2011). Mice received 20% (v/v) ethanol (EtOH; Koptec, Decon Labs, King of Prussia, PA) in tap water beginning 3 hr into the dark cycle, for 2 hr (i.e., 10:00 am to 12:00 pm) on three consecutive days. On the fourth day, EtOH was presented for 4 hr (i.e., 10:00 am to 2:00 pm), representing the “binge” day. Sipper tubes were weighed at the beginning and end of each drinking window. There were three days of abstinence between DID cycles. Mice underwent 4 cycles of DID, from postnatal day (PND) 28-52 or PND 30-54, during which control mice only had access to water but were otherwise treated identically (timeline in **Figure 1A**; ethanol consumption shown in **Figure 1B-F**). The small stagger in the first exposure day was to allow time-locked electrophysiology from littermate cohorts of mice. Blood ethanol concentrations (BECs) were examined in a pilot cohort of SST-IRES-Cre:Ai9 mice to validate the adolescent DID consumption model (**Figure 1D**). Tail blood samples were taken 30 min after the final binge session. BECs were determined using an Analox AM1 Analyzer (Analox Instruments, Lunenburg, MA). Additional cohorts of mice did not further undergo tail blood sampling to minimize stress effects.

**Figure 1.**
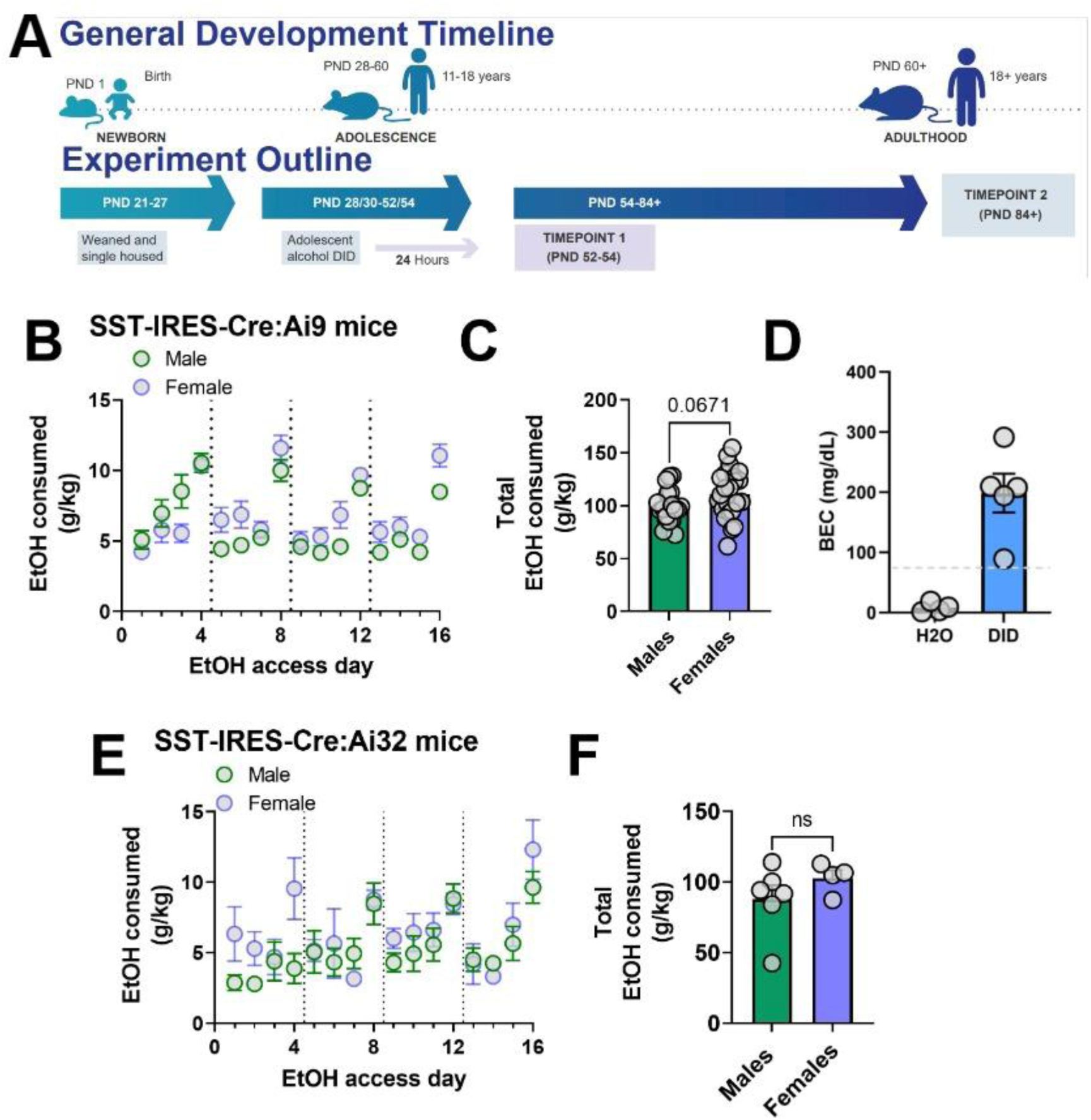
Overall experimental timeline and ethanol consumption during adolescent DID. (A) Adolescent development of mice versus humans. Alcohol is administered during PND 28/30-52/54, corresponding to a broad range of adolescence in humans. (B) Average daily ethanol consumption of male and female SST-IRES-Cre:Ai9 mice across the adolescent DID paradigm. (C) Total ethanol consumption from all 16 days of adolescent DID was not significantly different between male and female SST-IRES-Cre:Ai9 mice. (D) Blood ethanol concentrations from a pilot cohort of adolescent SST-IRES-Cre:Ai9 mice above the 80 mg / dL levels corresponding to a ‘binge’ episode. (E) Average daily ethanol consumption of male and female SST-IRES-Cre:Ai32 mice throughout adolescent DID exposure. (F) Total ethanol consumption across all days of adolescent DID was not significantly different between male and female SST-IRES-Cre:Ai32 mice.

### 2.3 Electrophysiology

Whole-cell current-clamp electrophysiology was performed as previously described (Dao et al., 2021, 2020). Electrophysiology was conducted at two separate timepoints to assess either the short-term (24 hr post-DID) or long-term (30 day post-DID) effects of adolescent alcohol on PL neuronal intrinsic excitability. Mice were anesthetized with inhaled isoflurane and rapidly decapitated. Brains were rapidly removed and immediately placed in ice-cold, oxygenated *N*-methyl-*D-*glucamine (NMDG) cutting solution containing the following, in mM: 93 NMDG, 2.5 KCl, 1.2 NaH2PO4, 30 NaHCO3, 20 HEPES, 25 dextrose, 5 ascorbic acid, 2 thiourea, 3 sodium pyruvate, 10 MgSO4-7 H2O, 0.5 CaCl2-2 H2O, 306-310 mOsm, pH to 7.4 (Ting et al., 2018). Coronal slices 300 µm thick containing the PL cortex were prepared using a Compresstome vibrating microtome (Precisionary Instruments). Slices recovered in heated (31◦C), oxygenated NMDG buffer for a maximum of eight minutes before resting in heated, oxygenated artificial cerebrospinal fluid (aCSF) containing in mM: 124 NaCl, 4.0 KCl, 1.2 MgSO4-7 H2O, 2.0 CaCl2-2H2O, 1 NaH2PO4-H2O, 305-308 mOsm, for at least 1 hr before recording. For experiments, slices were moved to a submerged chamber where they were consistently perfused with heated, oxygenated aCSF at a rate of 2 mL/min.

SST neurons in superficial layers of the PL cortex were identified based on the presence of the fluorescent tdTomato reporter using 565 nm LED excitation. Layer II/III pyramidal neurons were identified based on morphology, membrane characteristics (resistance and capacitance), and action potential width as previously characterized (Dao et al., 2021). Recording electrodes (3-6 MΩ) were pulled from thin-walled borosilicate glass capillaries with a Narishige P-100 Puller. Electrodes were filled with a potassium gluconate-based intracellular recording solution, containing in mM: 135 potassium gluconic acid, 5 NaCl, 2 MgCl2-6H2O, 10 HEPES, 0.6 EGTA, 4 Na2ATP, and 0.4 Na2-GTP, 287-290 mOsm, pH 7.35. Properties measured to assess the intrinsic excitability of SST and pyramidal neurons in the PL cortex included resting membrane potential (RMP), rheobase, action potential threshold, and the number of action potentials fired during a voltage-current (VI) protocol. In the VI protocol, increasing steps of depolarizing current were injected into the neuron (0-200 pA, increasing by 10 pA per step, each step lasting 300 ms). Negative current steps were included as a control. Experiments were performed at each neuron’s RMP and repeated holding the neurons at a common voltage of - 70mV.

SST-IRES-Cre:Ai32 mice were used to assess changes in local SST circuitry 24 hr following adolescent DID. Experiments to assess SST neuron-mediated input onto pyramidal and non-SST, non-pyramidal (putatively other GABAergic) neurons were conducted in voltage-clamp configuration, holding cells at –50mV, as described previously (Dao et al., 2021). To record optogenetically-evoked inhibitory postsynaptic currents, 3mM kynurenic acid was added to the recording ACSF to block glutamate receptors. Electrodes were filled with the potassium gluconate-based intracellular recording solution described above. Pyramidal and non-SST, non-pyramidal cells were identified based on morphology and membrane characteristics (membrane resistance and capacitance). Optogenetically-evoked inhibitory postsynaptic currents were elicited by 2, 1ms pulses of 470nm light delivered across an interstimulus interval ranging from 100-500 ms apart (Cool LED, Traverse City, MI, USA). Paired pulse ratio was calculated as the peak of the second optogenetically-evoked current divided by the first optogenetically-evoked current.

Signals were digitized at 10 kHz and filtered at 4 kHz using a MultiClamp 700B amplifier. Recordings were analyzed using Clampfit 10.7 software (Molecular Devices, Sunnyvale, CA, United States). A maximum of 2 cells per cell type (i.e. 2 SST and 2 pyramidal) were recorded from each mouse.

### 2.4 Analysis of Membrane Properties

Membrane properties were calculated using Clampfit 10.7 software (Molecular Devices, Sunnyvale, CA, United States). Interspike interval (ISI) was calculated by measuring the time between the peak of the first two action potentials of the 200 pA sweep during VI at RMP. Voltage sag ratio was calculated as the peak minimum voltage in the first 100ms of the most hyperpolarizing step of VI (-100pA) divided by the average voltage in the final 50ms of the step (Kinnischtzke et al., 2012). Input resistance was calculated as the slope of the I-V curve for hyperpolarizing current injection steps between -50pA and 0pA (Koppensteiner et al., 2019). The membrane time constant was determined by fitting an exponential curve to the end of the voltage response to a -50pA current injection step. Capacitance was calculated by dividing the time constant by the input resistance. Medium afterhyperpolarization (mAHP) was calculated using the 200pA injection step as the difference between the average membrane voltage prior to current injection and the peak minimum voltage in the 150ms after current injection. Action potential half-width was determined using the first action potential fired during rheobase at RMP.

### 2.5 Histology and SST cell counts

Twenty-four hours after the end of adolescent DID, mice were deeply anesthetized with Avertin (250 mg/kg) and perfused transcardially with phosphate buffered saline (PBS; pH 7.4) followed by 4% paraformaldehyde (PFA; pH 7.4). Brains were removed and postfixed in PFA overnight. Brains were sectioned at 40 µm using a Leica vibratome (VS 1200, Leica). Brain slices were placed in 1:10000 4′,6-diamidino-2-phenylindole (DAPI) for 10 minutes and washed with phosphate buffered saline (PBS; pH 7.4) on Fisherbrand™ Multi-Platform Shaker at 60 rpm 3x for 10 minutes. PL cortex containing sections were mounted on SuperFrost glass slides, air-dried, and coverslipped with ImmunoMount (Thermo Fisher Scientific, Waltham, MA, United States) mounting media. SST+ cell counts in SST-IRES-Cre:Ai9 mice, expressing the fluorophore tdTomato exclusively in SST-expressing neurons, were quantified using ImageJ (National Institutes of Health, Bethesda, MD, United States). The PL cortex was delineated and SST+ were automatically quantified under matched criteria for size, circularity, and intensity consistently with our previously published work (Dao et al., 2020; Suresh Nair et al., 2022). The threshold to highlight SST+ cells counted was set at 8.10±.01. Each ROI’s total SST cell count was divided by the ROI area to give a total SST+ density value (Smith et al., 2020). 7-8 slices were analyzed per mouse.

### 2.6 Data analysis, statistics, and figure preparation

Data were analyzed in GraphPad Prism 7.0 (San Diego, CA, United States). A 2-way ANOVA (factors: sex, adolescent DID condition) was used for SST cell counts and for membrane properties including RMP, rheobase, action potential threshold, and ISI. A mixed effects ANOVA (factors: sex, adolescent DID condition, current injection step) was used for VI experiments. When a main effect of sex or an interaction between sex and DID condition was seen, males and females were further analyzed separately by 2-way ANOVA. Tukey post-hoc tests were used when appropriate. Data are presented as the mean and standard error of the mean. One SST cell from the M H2O group was excluded from the ISI analysis at the 24 hr timepoint based on the ROUT test for outliers in GraphPad Prism. Excluding this cell did not impact the statistical outcome. One pyramidal cell from the M H2O group was excluded from all analysis on the 30-day electrophysiology timepoint because its rheobase and action potential threshold at RMP were identified as an outlier based on the ROUT test. Total alcohol consumption (g/kg) across all 4 cycles of adolescent DID was correlated with SST cell density and with the number of action potentials elicited by a 200 pA current injection step at RMP. Control mice were included in correlations with g/kg consumed = 0. Each correlation was reported as Pearson’s *r*.

## 3. RESULTS

### 3.1 Adolescent DID produces BECs above the threshold for binge drinking

Male and female SST-IRES-Cre:Ai9 and SST-IRES-Cre:Ai32 mice underwent a modified DID procedure spanning the bulk of the rodent adolescent window (**Figure 1A**). Daily ethanol consumption and total ethanol consumption for all SST-IRES-Cre:Ai9 (**Figure 1B-D**) and SST-IRES-Cre:Ai32 (**Figure 1E-F**) mice is shown below. “Binge” days consisting of 4 hr ethanol exposure occurred on sessions 4, 8, 12, and 16. Interestingly, we did not see a significant effect of sex on total ethanol consumption in SST-IRES-Cre:Ai9 mice who binged during adolescence (*t*(40) = 1.882, *p* = 0.0671). We also did not find a significant difference in the variance between males and females (*F*(22, 18) = 2.099, *p* = 0.1148). We further statistically confirmed that there were no unintended differences in ethanol consumption between the groups of SST-IRES-Cre:Ai9 mice used for both timepoints of electrophysiology experiments or cell counting experiments. A 2-way ANOVA (factors: sex, experiment) indicated no significant effect of sex or experiment (data not shown; F_sex_(1, 36) = 3.690, *p* = 0.0627; F_experiment_(2, 36) = 0.8641, *p* = 0.4300; F_sex x experiment_(2, 36) = 0.1433; *p* = 0.8670). Tail blood samples were taken from a pilot cohort of SST-IRES-Cre:Ai9 mice 30 minutes after the final binge session. All DID mice drank to BECs exceeding the 80 mg/dL threshold for binge drinking (**Figure 1D**). In addition, our DID procedure produces BECs comparable to previous studies using DID in adolescent C57 mice (Holstein et al., 2011; Wolstenholme et al., 2020) and matching our previous findings using adult mice (Dao et al., 2021). Due to the potential stress confound of tail blood sampling, subsequent cohorts of mice did not undergo BEC analysis, but g/kg of alcohol consumed was recorded every session. Daily (**Figure 1E**) and total (**Figure 1F**) ethanol consumption for SST-IRES-Cre:Ai32 mice used for electrophysiology experiments is shown below. There was no significant difference in alcohol consumption between male and female SST-IRES-Cre:Ai32 mice (*t*(8) = 1.147, *p* = 0.2845). This comparison shows that any differences in electrophysiology or cell counts were not because the mice used in these experiments inadvertently consumed different amounts of ethanol during adolescent DID.

### 3.2 SST neuronal excitability was increased 24 hours after drinking

We sought to characterize immediate changes in the intrinsic excitability of superficial PL SST neurons by performing electrophysiology 24 hr after the cessation of adolescent DID (**Figure 2**). RMP was not affected by sex or adolescent DID condition (representative traces in **Figure 2A-B,** RMP data in **Figure 2C;** F_sex_(1, 28) = 1.541, *p* = 0.2248; F_DID_(1, 28) = 3.380, *p* = 0.0766; F_sex x DID_(1, 28) = 0.09798; *p* = 0.7566). In rheobase experiments at RMP, we found a main effect of adolescent DID condition where SST neurons from alcohol exposed mice required significantly less current to fire an action potential (**Figure 2D;** F_DID_(1, 28) = 8.138, *p* = 0.0081), though there was no main effect of sex or interaction between sex and adolescent DID condition (F_sex_(1, 28) = 0.5451, *p* = 0.4665; F_sex x DID_(1, 28) = 0.02432; *p* = 0.8772). Action potential threshold was unaltered by adolescent DID condition (**Figure 2E;** F_sex_(1, 27) = 1.567, *p* = 0.2213; F_DID_(1, 27) = 0.6358, *p* = 0.4322; F_sex x DID_(1, 27) = 0.2807; *p* = 0.6006). In order to explore the potential underlying mechanisms of DID-induced changes in excitability, we also measured the interspike interval (ISI), or the interval between the first and second action potentials in the final step of the VI at RMP. Hyperpolarization-activated cyclic nucleotide (HCN) channels modulate properties related to neuronal excitability, including the ISI (Bohannon and Hablitz, 2018) and have previously been shown to be sensitive to adolescent alcohol in PL pyramidal neurons (Salling et al., 2018). However, the ISI in SST neurons was not altered by adolescent DID condition or sex (**Figure 2F;** F_sex_(1, 27) = 0.601, *p* = 0.4446; F_DID_(1, 27) = 2.225, *p* = 0.1474; F_sex x DID_(1, 27) = 0.886; *p* = 0.3547).

**Figure 2.**
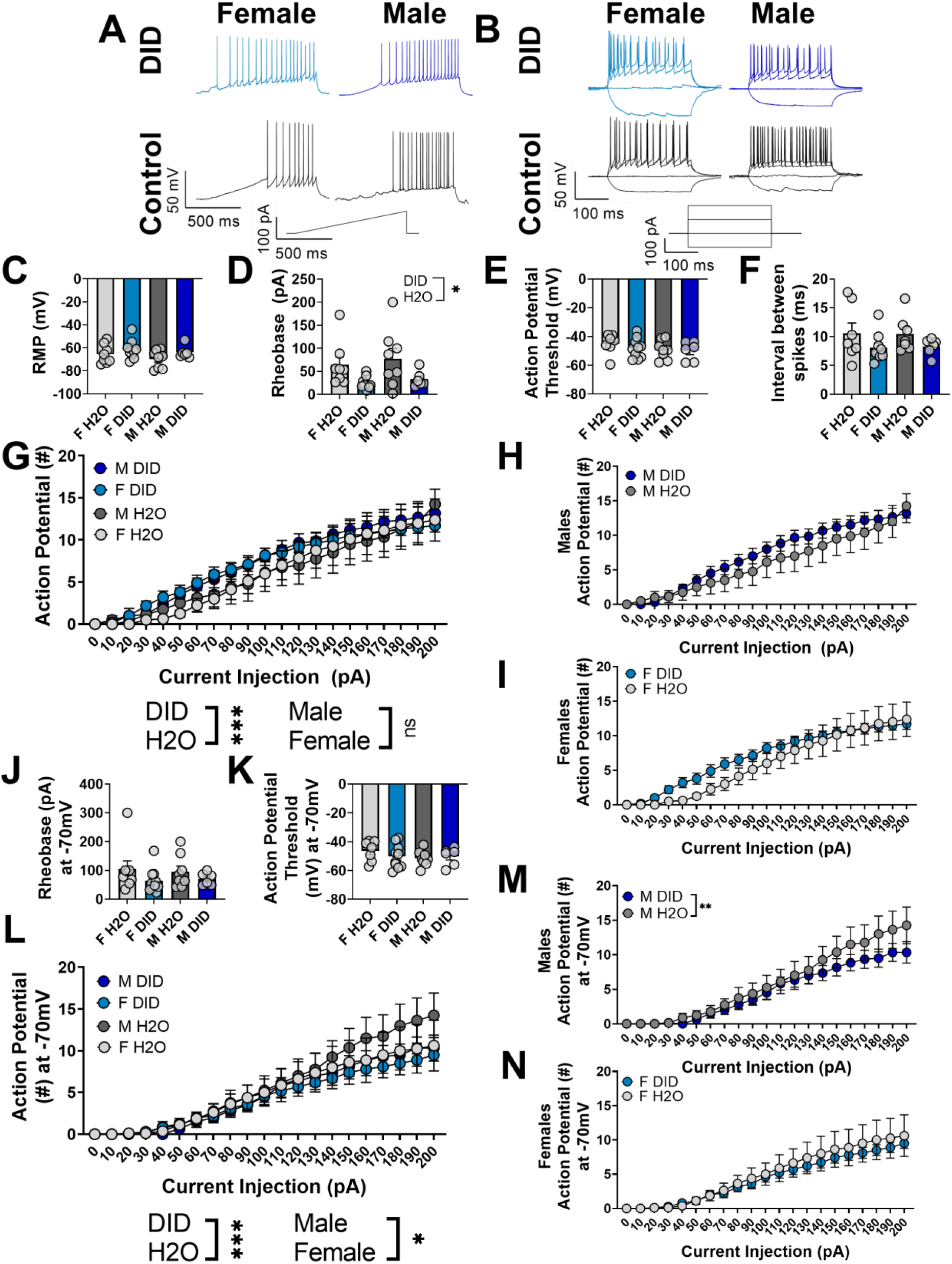
SST neurons are hyperexcitable 24 hr after adolescent DID. (A-B) Representative traces for rheobase and VI at RMP. (C-D) RMP is not altered by adolescent DID or sex, although SST neurons from mice which went through DID require less current to elicit an action potential. (E-F) Action potential threshold and interspike interval are not altered by adolescent DID or sex. (G) SST neurons from mice which drank alcohol during adolescence fired more action potentials compared to SST neurons from mice which did not drink during adolescence. (H-I) Data from (G), separated by sex for visualization. (J-K) Membrane properties of SST neurons were unaltered by sex or adolescent DID condition when neurons were held at -70mV. (L) There was a main effect of adolescent DID condition and of sex on the number of action potentials fired when SST neurons were held at -70mV. (M-N) Data from (L) split by sex for visualization. * indicates *p* < 0.05, ** indicates p < 0.01, *** indicates *p* < 0.0001. For panels A-L, *n* = 6 cells from 3 mice (M DID), 10 cells from 5 mice (F DID), 8 cells from 4 mice (M H2O), and 8 cells from 5 mice (F H2O).

We found a main effect of DID in the VI protocol at RMP (**Figure 2G;** females and males graphed separately in **Figure 2H-I**; mixed effects ANOVA with sex, alcohol exposure, and current injection as factors; F_DID_(1, 588) = 13.26, *p* = 0.0003). There was an expected effect of current injection (F_current_(20, 588) = 36.25, *p* < 0.0001). Higher current steps elicited more action potentials across all groups. There was no main effect of sex (F_sex_(1, 588) = 0.1753; *p* = 0.6756). All interactions were not significant (*p* > 0.05).

We repeated the experiments holding the SST neurons at the common membrane potential of -70 mV. At -70 mV, the rheobase was unaltered by sex or adolescent DID condition (**Figure 2J;** F_sex_(1, 28) = 0.003726, *p* = 0.9518; F_DID_(1, 28) = 2.525, *p* = 0.1233; F_sex x DID_(1, 28) = 0.1387; *p* = 0.7124). Action potential threshold was similarly not affected by sex or adolescent DID while neurons were held at -70 mV (**Figure 2K;** F_sex_(1, 27) = 1.003, *p* = 0.3256; F_DID_(1, 27) = 0.1473, *p* = 0.7041; F_sex x DID_(1, 27) = 0.8774; *p* = 0.3572). In the VI protocol at - 70 mV, we found significant main effects of sex (**Figure 2L**; mixed effects ANOVA with sex, alcohol exposure, and current injection as factors; F_sex_(1, 588) = 4.067, *p =* 0.0442) and adolescent DID condition (F_DID_(1, 588) = 9.437, *p* =0.0022). We found an expected main effect of current injection (F_current_(20, 588) = 32.23, *p* < 0.0001). Because of the main effect of sex, we further analyzed males and females separately using 2-way ANOVAs. In males, there was a main effect of adolescent DID (**Figure 2M**; F_DID_ (1, 252) = 7.003, *p* = 0.0087) and an expected main effect of current (F_current_(20, 252) = 16.35, *p* < 0.0001). When held at -70mV, SST neurons from male mice which went through adolescent DID fired fewer action potentials overall compared to male mice which did not drink alcohol. There was no significant interaction between current and DID (*p* > 0.05). In females, there was only a significant effect of current injection (**Figure 2N**; F_current_(20, 336) = 15.66, *p* < 0.0001) and no main effect of DID (F_DID_(20, 588) = 2.510, *p* = 0.1141) or interaction (*p* > 0.05). Together, this suggested modest increases in SST neuronal excitability in the PL cortex following adolescent DID.

We also assessed the membrane properties of SST neurons following adolescent DID (**Table 1**). There was no effect of adolescent DID or sex on input resistance of SST neurons (**Supplementary Figure 1A;** F_sex_(1, 27) = 1.463, *p* = 0.2370; F_DID_(1, 27) = 4.132, *p* = 0.0520; F_sex x DID_(1, 27) = 0.009791; *p* = 0.9219). There was a main effect of sex on membrane time constant (**Supplementary Figure 1B;** F_sex_(1, 27) = 12.66, *p* = 0.0014) but not DID condition (F_DID_(1, 27) = 0.056, *p* = 0.8142), nor any interaction (F_sex x DID_(1, 27) = 2.104, *p* = 0.1585). There was also a significant main effect of sex on membrane capacitance (**Supplementary Figure 1C;** F_sex_(1, 27) = 8.394, *p* = 0.0074), but not DID condition (F_DID_(1, 27) = 3.940, *p* = 0.0574) nor an interaction (F_sex x DID_(1, 27) = 3.138, *p* = 0.0878). Tukey post-hoc tests showed that the capacitance of SST neurons from female H2O-drinking mice was higher than the capacitance of SST neurons from all other groups. Voltage sag ratio, calculated from the most hyperpolarizing step (-100pA) of the VI protocol, was not altered by adolescent DID (**Supplementary Figure 1D;** F_DID_(1, 28) = 0.01522, *p* = 0.9027). However, there was a main effect of sex on the sag ratio (F_sex_(1, 28) = 8.324, *p* = 0.0074) with no significant interaction (F_sex x DID_(1, 28) = 2.281, *p* = 0.1421). Tukey post-hoc tests indicated that SST sag ratio was higher in H2O females than in H2O males. There was no significant effect of adolescent DID condition (**Supplementary Figure 1E;** F_DID_(1, 27) = 1.377, *p* = 0.2508) or sex (F_sex_(1, 27) = 0.2507, *p* = 0.6207), nor any interaction (F_sex x DID_(1, 27) = 0.05894, *p* = 0.8100), on mAHP in SST neurons. However, more SST neurons from adolescent DID mice exhibited no hyperpolarization following a 200pA current injection compared to SST neurons from control mice, shown by Fisher’s exact test (*p* = 0.0091). Finally, there was no difference in action potential half-width as a function of sex (**Supplementary Figure 1F;** F_sex_(1, 26) = 0.0008905, *p* = 0.9764) or DID (F_DID_(1, 26) = 0.5460, *p* = 0.4666), nor any interaction (F_sex x DID_(1, 26) = 0.8399, *p* = 0.3678).

**Table 1.** Membrane properties for SST neurons 24 hr following cessation of adolescent DID.

|  | F H2O | F DID | M H2O | M DID |
| --- | --- | --- | --- | --- |
| Input resistance (GΩ) | 0.3156 ± 0.04837 | 0.4123 ± 0.04239 | 0.2653 ± 0.03990 | 0.3529 ± 0.04693 |
| Membrane time constant (ms) | 37.50 ± 5.392 | 33.21 ± 2.143 | 19.76 ± 3.181 | 25.74 ± 3.337 |
| Membrane capacitance (pF) | 123.0 ± 13.04 | 86.00 ± 7.269 | 77.02 ± 9.725 | 74.91 ± 8.935 |
| Voltage sag ratio | 1.009 ± 0.004015 | 1.002 ± 0.003745 | 0.9888 ± 0.002655 | 0.9941 ± 0.008437 |
| mAHP (mV) | -1.962 ± 0.8335 | -0.2861 ± 0.1893 | -1.441 ± 0.6235 | -0.7873 ± 0.3697 |
| Action potential half width (ms) | 0.9259 ± 0.07039 | 0.9393 ± 0.04302 | 0.9927 ± 0.1118 | 0.8680 ± 0.03748 |

Data are presented as mean ± SEM.

### 3.3 SST excitability remained increased 30 days after drinking

Next, we investigated how long superficial SST neuronal hyperexcitability persisted after adolescent DID. In a separate cohort of mice, DID was repeated and then the mice were allowed to mature undisturbed from PND 54 to PND 84 before performing electrophysiology (**Figure 3**).

**Figure 3.**
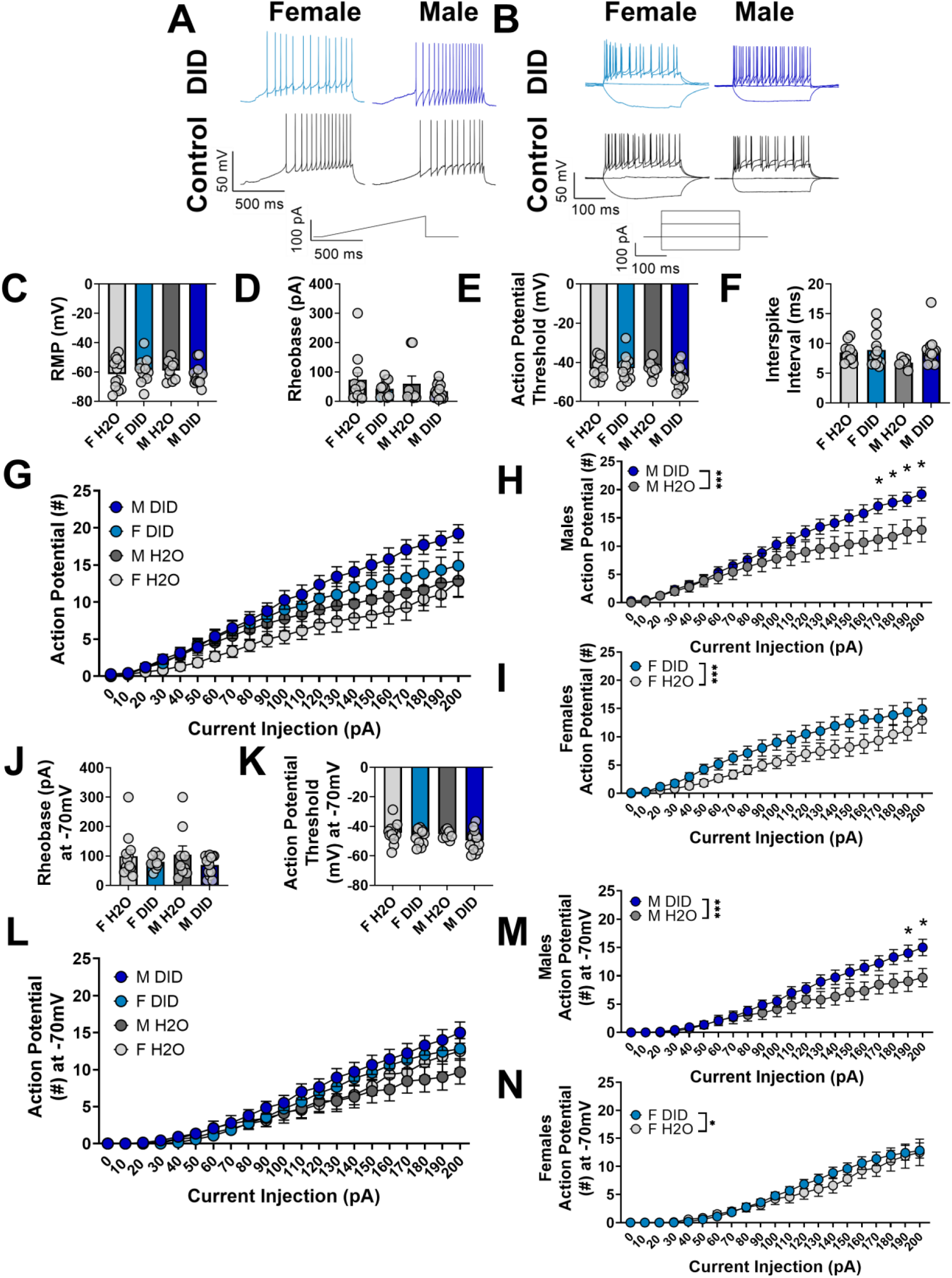
SST neurons remain hyperexcitable 30 days after cessation of adolescent DID. (A-B) Representative traces for rheobase and VI recordings. (C-F) Membrane properties and interspike interval of SST neurons were not altered as a function of adolescent DID or sex. (G) At RMP, we found a significant interaction of current injection step and adolescent DID condition, and a main effect of sex on current-induced action potential firing. (H-I) When the data from (G) at RMP were analyzed separately based on sex, there was a significant main effect of adolescent DID in both males and females. (J-K) Membrane properties of SST neurons were not altered when neurons were held at -70mV. (L) When held at -70mV, there was a significant interaction between sex and adolescent DID, as well as a main effect of current injection on current-induced action potential firing. (M-N) Data from (L) graphed separately by sex. When analyzed separately, there was a main effect of adolescent DID on the number of action potentials fired in both males and females. For panels A-L, *n* = 14 cells from 7 mice (M DID), 12 cells from 6 mice (F DID), 9 cells from 5 mice (M H2O), and 12 cells from 6 mice (F H2O). * indicates *p* < 0.05, *** indicates *p* < 0.0001. Main effects of adolescent DID or sex are indicated with brackets between group labels; significant differences in current-induced firing are indicated with * over specific current steps.

RMP was unaltered in SST neurons 30 days after adolescent DID (representative traces in **Figure 3A-B**, **Figure 3C** RMP data; F_sex_(1, 43) = 0.02353, *p* = 0.8788; F_DID_(1, 43) = 0.03061, *p* = 0.8619; F_sex x DID_(1, 43) = 1.298; *p* = 0.2609). Rheobase (**Figure 3D;** F_sex_(1, 43) = 0.3952, *p* = 0.5329; F_DID_(1, 43) = 2.871, *p* = 0.0974; F_sex x DID_(1, 43) = 0.04574; *p* = 0.8317) and action potential threshold (**Figure 3E;** F_sex_(1, 43) = 1.691, *p* = 0.2003; F_DID_(1, 43) = 1.361, *p* = 0.2498; F_sex x DID_(1, 43) = 1.536; *p* = 0.2220) were not changed as a function of sex or adolescent DID condition. There was no effect of adolescent DID condition or sex on the ISI (**Figure 3F**; F_sex_(1, 41) = 2.004, *p* = 0.1644; F_DID_(1, 41) = 2.903, *p* = 0.0960; F_sex x DID_(1, 43) = 1.098; *p* = 0.3009).

In the VI experiment at RMP, we found a significant interaction between current injection and DID condition (**Figure 3G**; mixed effects ANOVA with sex, alcohol exposure, and current injection as factors; F_DID x current_(20, 903) = 1.678; *p* = 0.0315) as well as a significant main effect of sex F_sex_(1, 903) = 30.89; *p* < 0.0001). All other interactions were not significant. Because we found a main effect of sex in our mixed effects analysis, we further analyzed the sexes separately as a 2-way ANOVA (factors: adolescent DID and current injection step). In males, we found significant main effects of current injection and adolescent DID condition (**Figure 3H;** F_current_(20, 441) = 31.25, *p* < 0.0001; F_DID_(1, 441) = 43.39, *p* < 0.0001; F_current x DID_(20, 43) = 1.502; *p* = 0.0758). In females, we found significant main effects of current and adolescent DID condition, with no significant interaction between adolescent DID and current injection (**Figure 3I;** F_current_(20, 462) = 24.02, *p* < 0.0001; F_DID_(1, 462) = 46.84, *p* < 0.0001; F_current x DID_(20, 43) = 0.5528; *p* = 0.9426).

We repeated these intrinsic excitability experiments holding the SST neurons at -70 mV. We found no main effect of sex or adolescent DID condition, or interaction between sex and adolescent DID condition, on rheobase at -70 mV (**Figure 3J;** F_sex_(1, 43) = 0.03137, *p* = .8602; F_DID_(1, 43) = 2.659, *p* = 0.1103; F_sex x DID_(1, 43) = 0.2176; *p* = 0.6433). Action potential threshold was also unaltered in SST neurons 30 days after adolescent DID (**Figure 3K;** F_sex_(1, 43) = 1.117, *p* = 0.2964; F_DID_(1, 43) = 3.395, *p* = 0.0723; F_sex x DID_(1, 43) = 0.3319; *p* = 0.5675). In the VI experiment conducted at -70 mV, we found a significant main effect of current (**Figure 3L;** mixed effects ANOVA with sex, alcohol exposure, and current injection as factors; F_current_(20, 903) = 67.44; *p* < 0.0001) and a significant interaction between adolescent DID condition and sex (F_sex x DID_(1, 903) = 8.528; *p* = 0.0036). Because of this interaction, we then analyzed the sexes separately using a 2-way ANOVA. In males, there was a significant main effect of current (**Figure 3M**; F_current_(20, 441) = 28.68; *p* < 0.0001) and of adolescent DID condition (F_DID_(1, 441) = 32.59; *p* < 0.0001) but no significant interaction (F_current x DID_(20, 462) = 1.452; *p* = 0.0940). In females, there was a significant expected main effect of current (**Figure 3N;** F_current_(20, 462) = 39.82; *p* < 0.0001) and a main effect of DID condition (F_DID_(1, 462) = 3.878; *p* = 0.0495) but no significant current x DID interaction (F_current x DID_(20, 462) = 0.3325; *p* = 0.9975). Together, this suggested that adolescent DID-induced changes in SST function persisted well after the cessation of binge drinking and are likely permanent adaptations.

We also assessed passive membrane properties of SST neurons, including input resistance, membrane capacitance, membrane time constant, voltage sag ratio, mAHP, and action potential half width (**Table 2; Supplementary Figure 2**). We found no effect of DID, or sex, or any interaction on the input resistance of SST neurons 30 days after the cessation of adolescent DID (**Supplementary Figure 2A;** F_sex_(1, 43) = 2.246, *p* = 0.1412; F_DID_(1, 43) = 0.001778, *p* = 0.9666; F_sex x DID_(1, 43) = 1.803; *p* = 0.1864). Membrane time constant was not altered by sex or DID condition (**Supplementary Figure 2B;** F_sex_(1, 42) = 0.001170, *p* = 0.9729; F_DID_(1, 42) = 0.04718, *p* = 0.8291; F_sex x DID_(1, 43) = 0.4830; *p* = 0.6709). Membrane capacitance was not significantly altered by sex or DID condition (**Supplementary Figure 2C;** F_sex_(1, 40) = 4.064, *p* = 0.0506; F_DID_(1, 40) = 3.982, *p* = 0.0528; F_sex x DID_(1, 40) = 0.01802; *p* = 0.8939). There was no effect of DID or sex, nor any interaction, on the voltage sag ratio of SST neurons 30 days after adolescent DID (**Supplementary Figure 2D;** F_sex_(1, 43) = 0.008184, *p* = 0.9283; F_DID_(1, 43) = 0.04718, *p* = 0.8291; F_sex x DID_(1, 43) = 0.4830; *p* = 0.6709). We found no effect of adolescent DID condition or sex on mAHP in SST neurons (**Supplementary Figure 2E;** F_sex_(1, 42) = 2.178, *p* = 0.2802; F_DID_(1, 42) = 0.9127, *p* = 0.3449; F_sex x DID_(1, 42) = 2.841; *p* = 0.0993). Fisher’s exact test revealed no difference in the number of SST cells which exhibited no mAHP between DID and H2O groups (*p* > 0.9999). Finally, neither sex nor adolescent DID condition affected the half-width of the first action potential in SST neurons (**Supplementary Figure 2F;** F_sex_(1, 40) = 0.02490, *p* = 0.8754; F_DID_(1, 40) = 0.04462, *p* = 0.8338; F_sex x DID_(1, 40) = 0.4503; *p* = 0.5061). These results indicate that membrane properties of SST neurons stabilize by 30 days after cessation of adolescent binge drinking, even though SST neurons remain hyperexcitable.

**Table 2.** Membrane properties for SST neurons assessed 30 days after end of adolescent DID.

|  | F H<sub>2</sub>O | F DID | M H<sub>2</sub>O | M DID |
| --- | --- | --- | --- | --- |
| Input resistance (GΩ) | 0.2833 ± 0.03825 | 0.3330 ± 0.04316 | 0.3920 ± 0.04874 | 0.3390 ± 0.02463 |
| Membrane time constant (ms) | 24.19 ± 3.595 | 28.22 ± 3.099 | 24.61 ± 4.603 | 28.05 ± 3.442 |
| Membrane capacitance (pF) | 75.23 ± 6.038 | 87.73 ± 9.211 | 60.80 ± 7.751 | 75.10 ± 3.429 |
| Voltage sag ratio | 0.9904 ± 0.02395 | 0.9863 ± 0.005058 | 0.9871 ± 0.007591 | 0.9884 ± 0.005815 |
| mAHP (mV) | -1.762 ± 0.3547 | -1.465 ± 0.4852 | -0.6322 ± 0.2317 | -1.706 ± 0.3762 |
| Action potential half width (ms) | 0.8850 ± 0.09236 | 0.8549 ± 0.03398 | 0.8308 ± 0.06914 | 0.8885 ± 0.05702 |

Data are presented as the mean ± SEM.

### 3.4 Adolescent DID does not change SST cell density in the PL cortex

We next assessed changes in PL SST cell count immediately following adolescent DID, as changes in the number of SST cells could impact the associated microcircuitry. Consistent with the first electrophysiology timepoint, mice were perfused 24 hr after the end of adolescent DID, with H2O-drinking controls perfused on the same day (representative images in **Figure 4A**). We found no effect of adolescent DID on the density of SST cells in the PL cortex (**Figure 4B**, F_DID_(1, 20) = 0.02833; *p* = 0.8680). There was no difference in SST cell density based on sex (F_sex_(1, 20) = 2.557; *p* = 0.1255) and there was no interaction between sex and adolescent DID condition (F_sex_(1, 20) = 0.03393; *p* = 0.8557).

**Figure 4.**
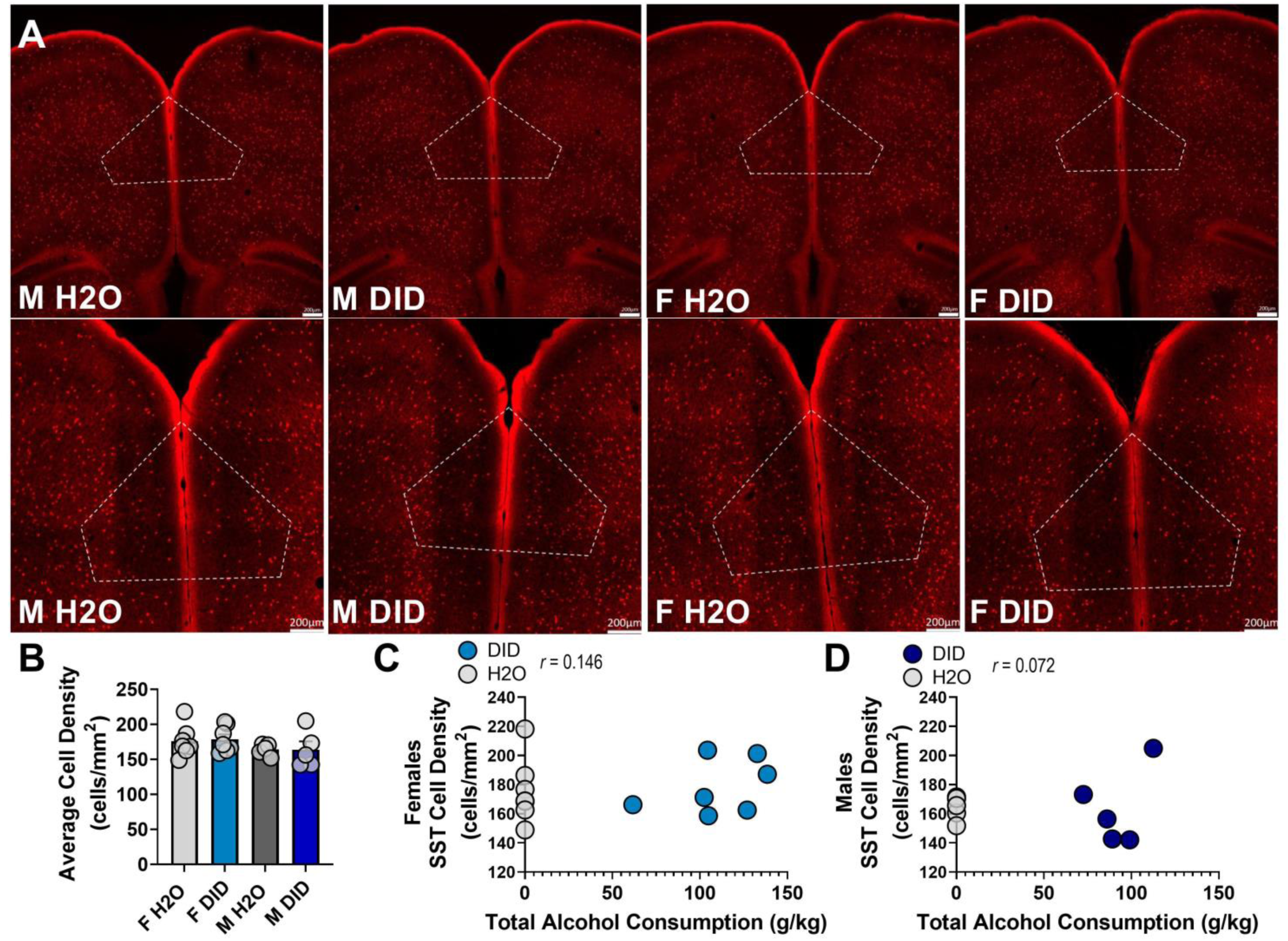
Adolescent DID does not alter the density of SST cells in the PL cortex. (A) Top row, representative images of SST cells in the PL cortex of SST-Ai9 mice. Scale bars in the bottom right of each image represent 200um. Bottom row, zoomed in images. Scale bars in the bottom right of each image represent 200 um. (B) There was no effect of sex or adolescent DID condition on SST cell density in the PL cortex. (C-D) Total alcohol consumption was not correlated with SST cell density in female or male mice. Filled circles represent individual DID mice; white circles represent individual H2O mice. For (B-D): *n* = 7 mice (F H2O and F DID), *n* = 5 mice (M H2O and M DID).

Next, we correlated individual levels of total alcohol consumption during adolescence with the number of PL SST cells to see if there was any association between drinking and SST cell density. There was no correlation between total alcohol consumption and SST cell density in females (**Figure 4C**, *r*(14) = 0.146, *p* = 0.619) or males (**Figure 4D**, *r*(10) = 0.072, *p* = 0.844). Together, these results indicate that adolescent binge drinking does not affect the density of SST cells in the PL cortex.

### 3.5 Pyramidal neuron excitability is reduced 24 hours after drinking

To determine if there are compensatory circuit changes after adolescent DID, we measured the intrinsic excitability of layer II/III PL pyramidal neurons 24 hr after binge drinking ended (**Figure 5**). We found no effect of adolescent DID condition or sex on RMP (representative traces for rheobase and VI in **Figure 5A-B**, RMP data in **Figure 5C**; F_sex_(1, 30) < 0.0001; *p* = 0.9954; F_DID_(1, 30) = 1.145; *p* = 0.2932; F_sex x DID_(1, 30) = 0.2332; *p* = 0.6327). Rheobase was also unaltered by sex and adolescent DID condition (**Figure 5D**; F_sex_(1, 30) = 0.892; *p* = 0.3522; F_DID_(1, 30) = 3.498; *p* = 0.0712; F_sex x DID_(1, 30) = 0.746; *p* = 0.3946), as was action potential threshold (**Figure 5E**; F_sex_(1, 30) = 0.4251; *p* = 0.5194; F_DID_(1, 30) = 0.05552; *p* = 0.8153; F_sex x DID_(1, 30) = 1.037; *p* = 0.3167). However, there was a significant interaction between current injection step and DID condition during the VI protocol (**Figure 5F**; mixed effects ANOVA with sex, alcohol exposure, and current injection as factors; F_current x DID_(20, 630) = 1.859; *p* = 0.0129), as well as an interaction between sex and adolescent DID condition (F_sex x DID_(1, 630) = 4.008; *p* = 0.0457), on current-induced action potential firing. Because of the interaction between sex and adolescent DID, we analyzed the data separately by sex. A 2-way ANOVA revealed a main effect of current injection step and of adolescent DID condition in males (**Figure 5G**; F_current_(20, 294) = 7.633; *p* < 0.0001, F_DID_(1, 294) = 31.63; *p* < 0.0001; F_current x DID_(20, 294) = 1.378; *p* = 0.1312). Similarly, there were significant main effects of current injection and adolescent DID condition in pyramidal neurons from females (**Figure 5H**; F_current_(20, 336) = 14.93; *p* < 0.0001, F_DID_(1, 336) = 16.21; *p* < 0.0001; F_current x DID_(20, 336) = 0.5181; *p* = 0.9587). In both sexes, PL pyramidal neurons showed reduced excitability following adolescent DID.

**Figure 5.**
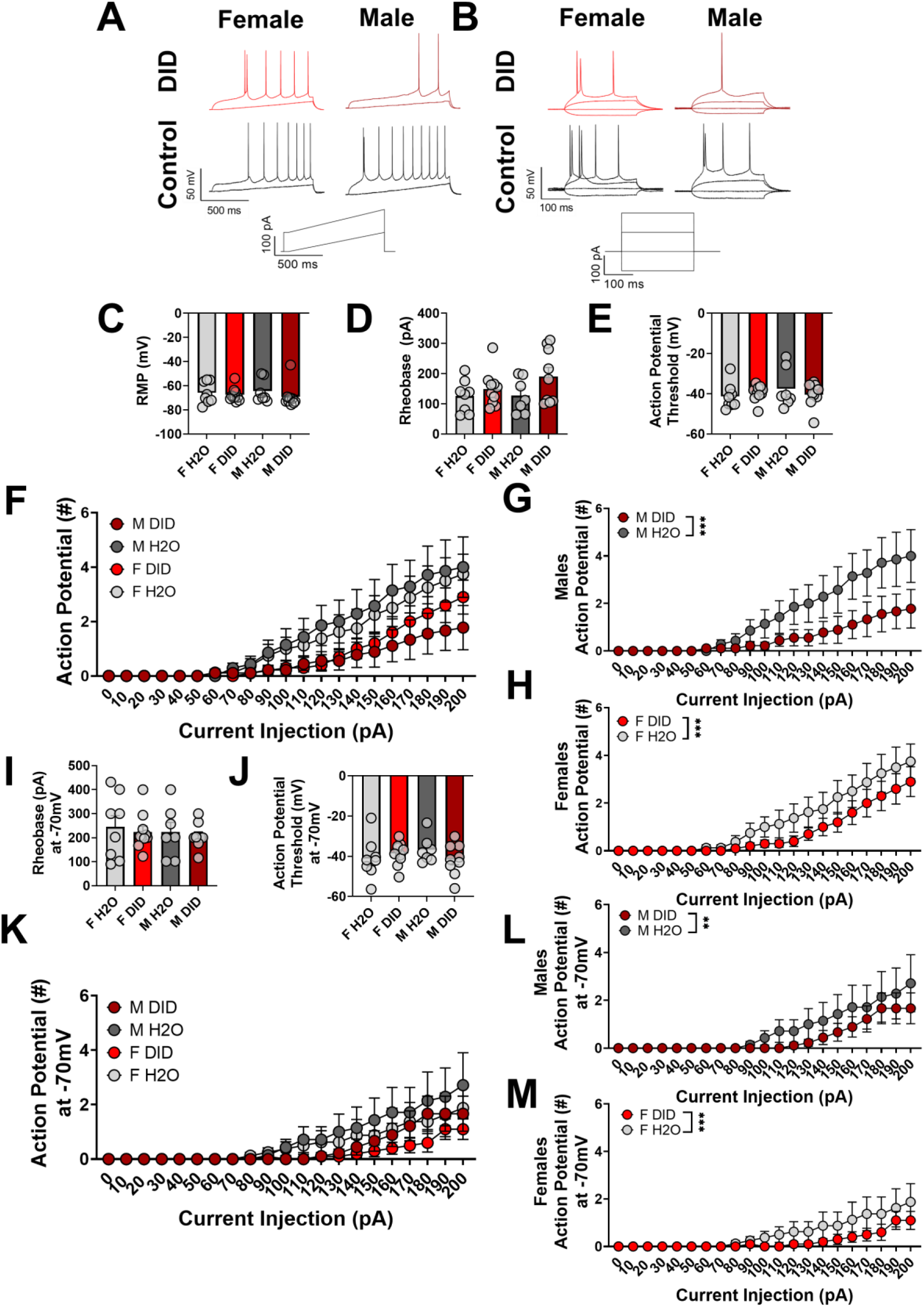
Pyramidal neurons are hypoexcitable 24 hr after adolescent DID. (A-B) Representative traces for rheobase and VI experiments in pyramidal neurons 24 hr after binge drinking. (C-E) There were no significant differences in RMP, rheobase, or action potential threshold as a function of adolescent DID or sex. (F) At RMP, we found a significant interaction between sex and adolescent DID as well as an expected main effect of current on current-induced action potential firing. (G-H) Data from (F) graphed separately by sex. When analyzed separately by sex, adolescent DID reduced action potential firing in both males and females. (I-J) Rheobase and action potential threshold were unaltered by adolescent DID condition or sex when neurons were held at -70mV. (K) At -70mV, there were significant main effects of sex, adolescent DID, and current injection step on action potential firing. (L-M) Data from (K) graphed separately by sex. Pyramidal neurons from male and female mice which underwent adolescent DID fired fewer action potentials when neurons were held at - 70mV. ** indicates p < 0.01, *** indicates *p* < 0.0001. For panels (A-M), *n* = 9 cells from 5 mice (M DID), 10 cells from 5 mice (F DID), 7 cells from 4 mice (M H2O), and 8 cells from 4 mice (F H2O).

We then assessed the excitability of pyramidal neurons while holding the neurons at a common voltage of -70 mV. There was no effect of sex or adolescent DID condition on rheobase in neurons held at -70 mV (**Figure 5I**; F_sex_(1, 30) = 0.06683; *p* = 0.7978, F_DID_(1, 30) = 0.1914; *p* = 0.6649; F_sex x DID_(1, 30) = 1.880; *p* = 0.1804). Neither sex nor adolescent DID condition altered action potential threshold when the neurons were held at - 70mV (**Figure 5J**; F_sex_(1, 30) = 0.3422; *p* = 0.5630, F_DID_(1, 30) = 0.3345; *p* = 0.5674; F_sex x DID_(1, 30) = 0.005682; *p* = 0.9404). We found significant main effects of current injection, sex, and adolescent DID on current-induced action potential firing when pyramidal neurons were held at -70mV (**Figure 5K**; mixed effects ANOVA with sex, alcohol exposure, and current injection as factors; F_current_(20, 630) = 10.57; *p* < 0.0001; F_sex_(1, 630) = 6.202; *p* = 0.0130, F_DID_(1, 630) = 18.26; *p* < 0.0001). All interactions were not significant (*p* > 0.05). Because we found a main effect of sex, we analyzed the data from males and females separately using a 2-way ANOVA. In males, there was an expected main effect of current injection (**Figure 5L**; F_current_(20, 294) = 5.701; *p* < 0.0001) and a main effect of adolescent DID (F_DID_(1, 294) = 6.874; *p* = 0.0092), but no interaction between current and DID (F_current x DID_(20, 294) = 0.3311; *p* = 0.9974). In females, we found similar effects of current injection (**Figure 5M**; F_current_(20, 336) = 4.724; *p* < 0.0001) and adolescent DID (F_DID_(1, 336) = 12.86; *p* = 0.0004) with no interaction (F_current x DID_(20, 336) = 0.5633; *p* = 0.9359). Together these results indicate an immediate reduction in pyramidal neuron excitability in both males and females after adolescent binge drinking.

We measured the membrane properties of pyramidal neurons 24 hr after adolescent DID (**Table 3, Supplementary Figure 3**). We found no effect of sex or adolescent DID on input resistance (**Supplementary Figure 3A,** F_sex_(1, 30) = 0.5964; *p* = 0.4460, F_DID_(1, 30) = 0.3542; *p* = 0.5562; F_sex x DID_(1, 30) = 1.141; *p* = 0.2939), membrane time constant (**Supplementary Figure 3B,** F_sex_(1, 30) = 0.004752; *p* = 0.9455, F_DID_(1, 30) = 0.4911; *p* = 0.4888; F_sex x DID_(1, 30) = 0.3414; *p* = 0.5634), or membrane capacitance (**Supplementary Figure 3C,** F_sex_(1, 30) = 0.2024; *p* = 0.6560, F_DID_(1, 30) = 0.05060; *p* = 0.8235; F_sex x DID_(1, 30) = 0.2706; *p* = 0.6067). There was a significant main effect of adolescent DID condition on voltage sag ratio (**Supplementary Figure 3D,** F_sex_(1, 30) = 4.635; *p* = 0.0395), but no effect of sex (F_sex_(1, 30) = 1.514; *p* = 0.2281) nor any interaction (F_sex x DID_(1, 30) = 0.0002474; *p* = 0.9876). There was no effect of sex (F_sex_(1, 30) = 3.710; *p* = 0.0636), adolescent DID (F_DID_(1, 30) = 0.5735; *p* = 0.4548), nor any interaction (F_sex x DID_(1, 30) = 0.01474; *p* = 0.9042) on mAHP (**Supplementary Figure 3E)**. Half-width of the first action potential fired in rheobase was also unaltered by sex or adolescent DID condition (**Supplementary Figure 3F,** F_sex_(1, 30) = 0.004124; *p* = 0.9493, F_DID_(1, 30) = 0.7009; *p* = 0.4096; F_sex x DID_(1, 30) = 0.1210; *p* = 0.7306).

**Table 3.** Membrane properties of pyramidal neurons 24 hr after cessation of adolescent DID.

|  | F H2O | F DID | M H2O | M DID |
| --- | --- | --- | --- | --- |
| Input resistance (GΩ) | 0.1242 ± 0.01499 | 0.1317 ± 0.01317 | 0.1534 ± 0.01892 | 0.1270 ± 0.01648 |
| Membrane time constant (ms) | 13.19 ± 1.236 | 12.95 ± 2.157 | 14.58 ± 2.595 | 11.85 ± 2.164 |
| Membrane capacitance (pF) | 112.5 ± 11.58 | 100.1 ± 13.58 | 96.41 ± 14.48 | 101.3 ± 22.26 |
| Voltage sag ratio | 0.9910 ± 0.002267 | 0.9944 ± 0.001540 | 0.9930 ± 0.001274 | 0.9964 ± 0.000937 |
| mAHP (mV) | -1.160 ± 0.3482 | -0.9892 ± 0.3193 | -0.6756 ± 0.1746 | -0.4399 ± 0.1042 |
| Action potential half width (ms) | 1.599 ± 0.08680 | 1.536 ± 0.1492 | 1.636 ± 0.1319 | 1.484 ± 0.09931 |

Data presented as the mean ± SEM

### 3.6 Pyramidal neuron excitability 30 days after drinking

We next measured persistent changes in layer II/III pyramidal neuron intrinsic excitability 30 days after adolescent DID concluded (**Figure 6**). Neither adolescent DID condition nor sex altered RMP of pyramidal neurons (representative traces for rheobase and VI in **Figure 6A-B,** RMP data in **Figure 6C;** F_sex_(1, 38) = 0.8349; *p* = 0.3666; F_DID_(1, 38) = 1.178; *p* = 0.2847; F_sex x DID_(1, 38) = 0.07644; *p* = 0.7837). Rheobase did not change as a function of sex or adolescent DID condition (**Figure 6D;** F_sex_(1, 38) = 0.9305; *p* = 0.3408; F_DID_(1, 38) = 0.4772; *p* = 0.4939; F_sex x DID_(1, 38) = 0.8061; *p* = 0.3749). Action potential threshold was also unaltered (**Figure 6E;** F_sex_(1, 38) = 9.480 E-6; *p* = 0.9976; F_DID_(1, 38) = 0.006398; *p* = 0.9367; F_sex x DID_(1, 39) = 0.9443; *p* = 0.3373). In the VI protocol at RMP, we found an expected main effect of current (**Figure 6F;** mixed effects ANOVA with sex, alcohol exposure, and current injection as factors; F_current_(20, 798) = 18.68; *p <* 0.0001) and a main effect of sex (F_sex_(1, 798) = 5.129; *p* = 0.0238) but not an effect of adolescent DID condition (F_DID_(1, 798) = 0.3502; *p* = 0.5542). There was a significant interaction between sex and adolescent DID condition (F_sex x DID_(1, 798) = 5.902; *p* = 0.0153). In males, there was an expected effect of current injection (**Figure 6G;** F_current_(20, 315) = 8.125; *p* < 0.0001) but no effect of adolescent DID condition (F_DID_(1, 315) = 1.174; *p* = 0.2794) and no interaction (F_current x DID_(20, 315) = 0.04198; *p* > 0.9999). However, in females, there were significant main effects of current (**Figure 6H;** F_current_(20, 483) = 10.46; *p* > 0.0001) and adolescent DID condition (F_DID_(1, 483) = 6.525; *p* = 0.0109), but no interaction (F_current_ x DID(20, 483) = 0.5006; *p* = 0.9664).

**Figure 6.**
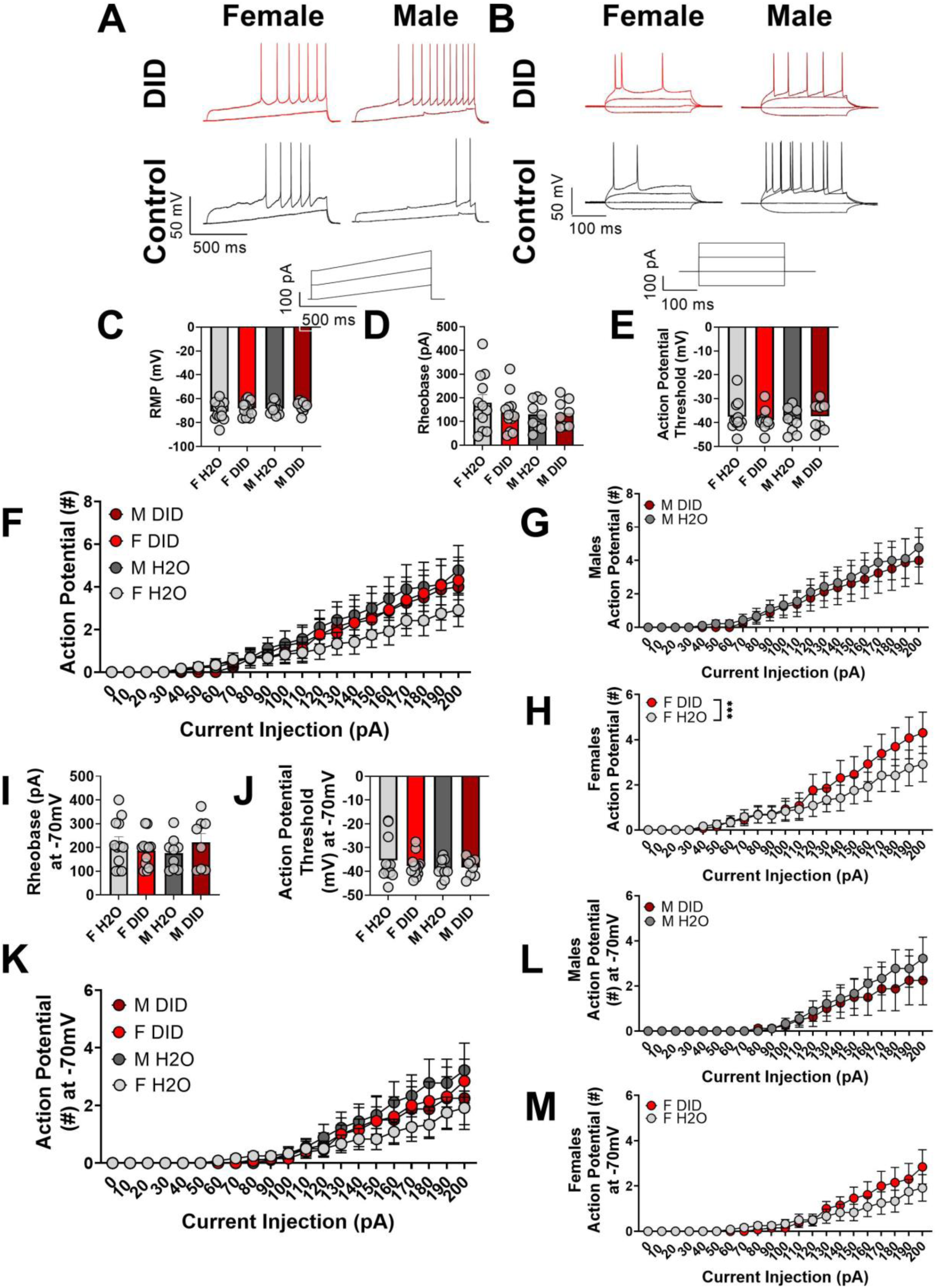
Pyramidal neuron excitability is not altered 30 days after adolescent DID exposure. (A-B) Representative traces for rheobase and VI experiments. (C-E) Pyramidal neuron RMP, rheobase, and action potential threshold are unaltered by adolescent DID and sex. (F) Current-induced action potential firing is not altered in pyramidal neurons by DID or sex 30 days after the cessation of adolescent DID. (G-H) Data from (F) plotted separately by sex for visualization. (I-J) Pyramidal neuron membrane properties are unaltered when held at -70mV. (K) Current-induced firing is not altered by adolescent DID or sex when neurons are held at -70mV. (L-M) Data from (K) graphed separated by sex for visualization. For panels A-L, *n =* 8 cells from 4 mice (M DID), 13 cells from 8 mice (F DID), 9 cells from 5 mice (M H2O), and 12 cells from 6 mice (F H2O).

We repeated the experiments holding the neurons at the common membrane potential of -70 mV. At -70 mV, there was no effect of adolescent DID condition or sex on rheobase (**Figure 6I;** F_sex_(1, 38) = 0.01121; *p* = 0.9162; F_DID_(1, 38) = 0.07281; *p* = 0.7887; F_sex x DID_(1, 38) = 1.990; *p* = 0.1665) or action potential threshold (**Figure 6J;** F_sex_(1, 38) = 1.217; *p* = 0.2770; F_DID_(1, 38) = 0.1931; *p* = 0.6628; F_sex x DID_(1, 38) = 0.6183; *p* = 0.4366). In the VI protocol at -70 mV, we found an expected main effect of current (**Figure 6K;** mixed effects ANOVA with sex, alcohol exposure, and current injection as factors; F_current_(20, 798) = 17.44; *p <* 0.0001). However, there was no main effect of sex (F_sex_(1, 798) = 3.384; *p =* 0.0662) or adolescent DID condition (F_DID_(1, 798) = 0.006136; *p* = 0.9376). There was a significant interaction between sex and adolescent DID condition (F_sex x DID_(1, 798) = 4.734; *p* = 0.0299). As above, we further analyzed the sexes separately as a 2-way ANOVA due to the sex differences seen at the 24 hr timepoint and throughout the circuit. We found an expected main effect of current injection, but no effect of adolescent DID in males (**Figure 6L;** F_current_ (20, 315) = 7.164; *p <* 0.0001; F_DID_ (1, 315) = 1.658; *p* = 0.1988) or in females (**Figure 6M;** F_current_ (20, 483) = 10.29; *p* < 0.0001; F_DID_ (1, 483) = 3.343; *p* = 0.0681). Together, these results suggest that PL pyramidal neurons in layer II/III may rebound towards hyperexcitability 30 days after adolescent DID in females, potentially to compensate for permanent increases in GABAergic signaling arising from overactive SST neurons.

We also analyzed the membrane properties of PL pyramidal neurons 30 days after adolescent DID exposure ended (**Table 4; Supplementary Figure 4**). Pyramidal neuron membrane resistance was not altered as a function of adolescent DID condition or sex (**Supplementary Figure 4A;** F_sex_(1, 38) = 0.1019; *p* = 0.7513; F_DID_(1, 38) = 3.616 E-5; *p* = 0.9952; F_sex x DID_(1, 38) = 0.07837; *p* = 0.7810). There was no effect of sex or adolescent DID condition on membrane time constant (**Supplementary Figure 4B;** F_sex_(1, 38) = 0.0008447; *p* = 0.9770; F_DID_(1, 38) = 0.1189; *p* = 0.7321; F_sex x DID_(1, 38) = 0.01308; *p* = 0.9095) or membrane capacitance (**Supplementary Figure 4C;** F_sex_(1, 38) = 0.7358; *p* = 0.3964; F_DID_(1, 38) = 0.6178; *p* = 0.4367; F_sex x DID_(1, 38) = 0.2158; *p* = 0.6449). We found a significant interaction between adolescent DID condition and sex in the voltage sag ratio test (**Supplementary Figure 4D;** F_sex x DID_(1, 38) = 4.387; *p* = 0.0429) but no significant main effects (F_sex_(1, 38) = 0.2018; *p* = 0.6558; F_DID_(1, 38) = 0.08552; *p* = 0.7715). There was no effect of adolescent DID condition or sex on action potential half-width (**Supplementary Figure 4E;** F_sex_(1, 35) = 1.982; *p* = 0.1680; F_DID_(1, 35) = 1.835; *p* = 0.1842; F_sex x DID_(1, 35) = 0.002498; *p* = 0.9604) or medium afterhyperpolarization (**Supplementary Figure 4F;** F_sex_(1, 38) = 0.3252; *p* = 0.5719; F_DID_(1, 38) = 0.05950; *p* = 0.8086; F_sex x DID_(1, 38) = 0.01810; *p* = 0.8937).

**Table 4.** Membrane properties of PL pyramidal neurons 30 days after adolescent DID.

|  | F H2O | F DID | M H2O | M DID |
| --- | --- | --- | --- | --- |
| Input resistance (GΩ) | 0.15270 ± 0.02783 | 0.1464 ± 0.01675 | 0.1536 ± 0.01741 | 0.1601 ± 0.02403 |
| Membrane time constant (ms) | 15.13 ± 2.486 | 15.63 ± 1.583 | 14.94 ± 2.212 | 15.94 ± 2.197 |
| Membrane capacitance (pF) | 113.9 ± 14.78 | 118.6 ± 13.76 | 95.00 ± 7.325 | 112.9 ± 17.17 |
| Voltage sag ratio | 0.9922 ± 0.001347 | 0.9957 ± 0.001132 | 0.9946 ± 0.0008865 | 0.9920 ± 0.002367 |
| mAHP (mV) | -0.7991 ± 0.3279 | -0.7692 ± 0.1574 | -0.9916 ± 0.3347 | -0.8882 ± 0.2110 |
| Action potential half width (ms) | 1.497 ± 0.1055 | 1.347 ± 0.08472 | 1.342 ± 0.08158 | 1.203 ± 0.1552 |

Data presented as mean ± SEM

### 3.7 Optogenetically-evoked SST neuron GABA release onto pyramidal and non-pyramidal populations immediately following adolescent DID

Our intrinsic excitability data suggests broad changes in SST and pyramidal neuron activity following adolescent exposure to alcohol consumption. However, it is unclear how SST neuronal changes may drive changes in the PL microcircuit. We have previously characterized PL SST neuronal GABA release onto both pyramidal and non-pyramidal populations (Dao et al., 2021). Here, we used a line of mice expressing ChR2 specifically in SST neurons (SST-IRES-Cre:Ai32). 24 hrs following adolescent DID, slices were prepared from male and female mice, and we recorded optically-evoked IPSCs onto pyramidal and non-pyramidal, non-SST neurons (**see Figure 7A** for schematic and **Figure 7B for** representative traces). Paired pulse ratio (PPR) was assessed at interstimulus intervals of 100, 250, and 500 msec. Broadly speaking, we did not see significant changes in PPR following adolescent DID. PPR of optogenetically-evoked SST GABA transmission onto pyramidal neurons was not changed at 100 msec (**Figure 7C;** F_sex_(1, 35) = 1.366; *p* = 0.2504; F_DID_(1, 35) = 2.055; *p* = 0.1606; F_sex x DID_(1, 35) = 0.2956; *p* = 0.5901), 250 msec (**Figure 7D;** F_sex_(1, 35) = 0.2037; *p* = 0.6545; F_DID_(1, 35) = 0.9200; *p* = 0.3441; F_sex x DID_(1, 35) = 0.1929; *p* = 0.6632), or 500 msec (**Figure 7E**); F_sex_(1, 35) = 1.075; *p* = 0.3068; F_DID_(1, 35) = 0.001251; *p* = 0.9720; F_sex x DID_(1, 35) = 0.2004; *p* = 0.6571) interstimulus intervals. Similarly, PPR of optogenetically-evoked SST GABA transmission onto non-pyramidal neurons was not changed at these intervals (100 msec data shown in **Figure 7F**; F_sex_(1, 29) = 1.208; *p* = 0.2808; F_DID_(1, 29) = 0.4023; *p* = 0.5309; F_sex x DID_(1, 29) = 0.5178; *p* = 0.4775), 250 msec shown in **Figure 7G**; F_sex_(1, 29) = 0.07830; *p* = 0.7816; F_DID_(1, 29) = 0.07095; *p* = 0.7918; F_sex x DID_(1, 29) = 1.882; *p* = 0.1807), or 500 msec shown in **Figure 7H;** F_sex_(1, 29) = 0.7940; *p* = 0.3802; F_DID_(1, 29) = 0.005444; *p* = 0.9417; F_sex x DID_(1, 29) = 1.080; *p* = 0.3073). We also compared the area-under-the-curve (AUC) across these three stimulus intervals. Comparison of AUCs for females (**Figure 7I**) and males (**Figure 7J**) showed no effect of adolescent DID on the AUC pyramidal neurons from females *(t*(13) = 0.3840; *p* = 0.7072) or males (*t*(14) = 0.4101; *p = 0.6879*), or on the AUC of non-pyramidal neurons from females (*t*(9) = 0.6479; *p* = 0.5332) or males (*t*(12) = 0.4001; *p* = 0.6961).

**Figure 7.**
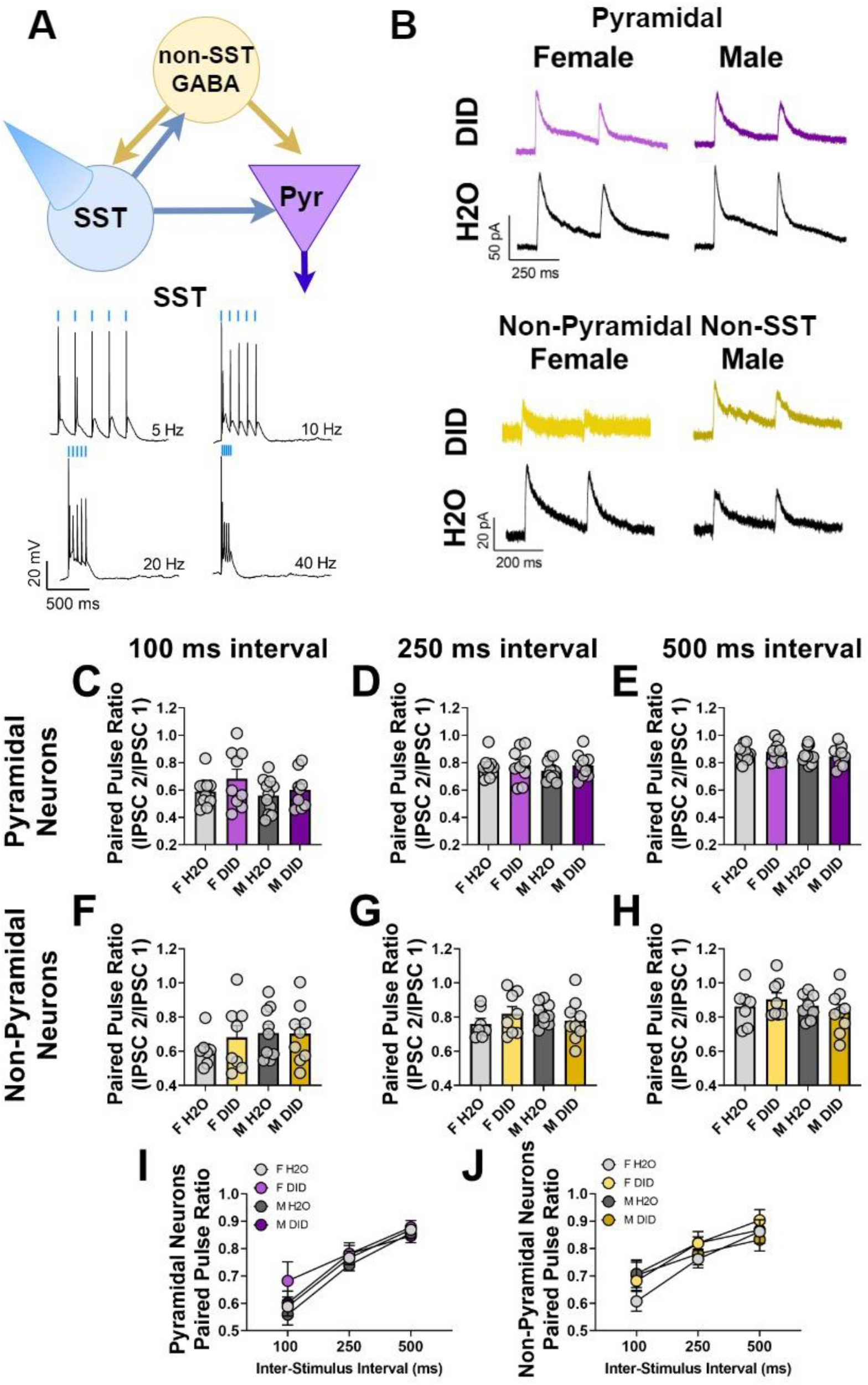
Optogenetically-evoked SST GABA release onto pyramidal neurons and non-pyramidal neurons is not altered 24 hrs after adolescent DID. A, schematic of experimental prep and representative optogenetically-evoked action potentials from SST cells at 5-40 hz. ChR2-mediated GABA release was optogenetically evoked from SST neurons. B, representative traces of PPR experiments in female (left) and male (right), DID and H20 mice, onto both pyramidal (top; scale bar represents 5o pA by 250 msec) and non-pyramidal (bottom; scale bar represents 20 pA by 200 msec) cells. (C-E) There was no significant difference in the paired pulse ratio of optogenetically-evoked inhibitory currents onto pyramidal neurons 24 hr after adolescent DID. (F-H) Paired pulse ratio of optogenetically-evoked inhibitory currents onto non-SST, non-pyramidal neurons was not altered 24 hr after adolescent DID. (I-J) AUC of paired pulse ratio across all 3 inter-stimulus intervals was not affected by sex or by adolescent DID condition.

### 3.8 Relationship between altered excitability and alcohol consumed during the binge paradigm

We next sought to understand how individual variability in alcohol consumption drove overall changes in cortical excitability. As the most prominent changes were identified in the VI protocol, we correlated the number of action potentials fired during the last step of the VI in SST neurons and pyramidal neurons with the g/kg alcohol consumed throughout the 4 week DID exposure (**Figure 8**). Total alcohol consumption was not correlated with SST action potential firing 24 hr after DID in males (**Figure 8A**; *r*(12) = -0.125, *p* = 0.669) or females (**Figure 8B**; *r*(16) = -0.044, *p* = 0.863). In SST neurons from adult male mice (30 days after the cessation of DID), total alcohol consumption across the adolescent DID paradigm was significantly correlated with the number of action potentials fired during the last step of VI (**Figure 8C**; *r*(21) = 0.464, *p* = 0.026). However, total drinking was not correlated with action potential firing in adult females (**Figure 8D;** *r*(22) = 0.137, *p* = 0.523). In pyramidal neurons recorded 24 hr after binge drinking, total alcohol consumption during the 4 weeks of adolescent DID was not significantly correlated with action potentials fired in males (**Figure 8E**; *r*(14) = -0.421; *p* = 0.105) or females (**Figure 8F;** *r*(16) = -0.282, *p* = 0.256). In pyramidal neurons recorded 30 days after adolescent DID, total alcohol drinking during adolescence was not correlated with current-induced action potential firing in males (**Figure 8G**; *r*(16) = -0.042, *p* = 0.870) or females (**Figure 8H;** *r*(23) = 0.250, *p* = 0.229). Together, these results indicate that alcohol consumption in adolescence is correlated with SST neuronal dysfunction in adult male mice.

**Figure 8.**
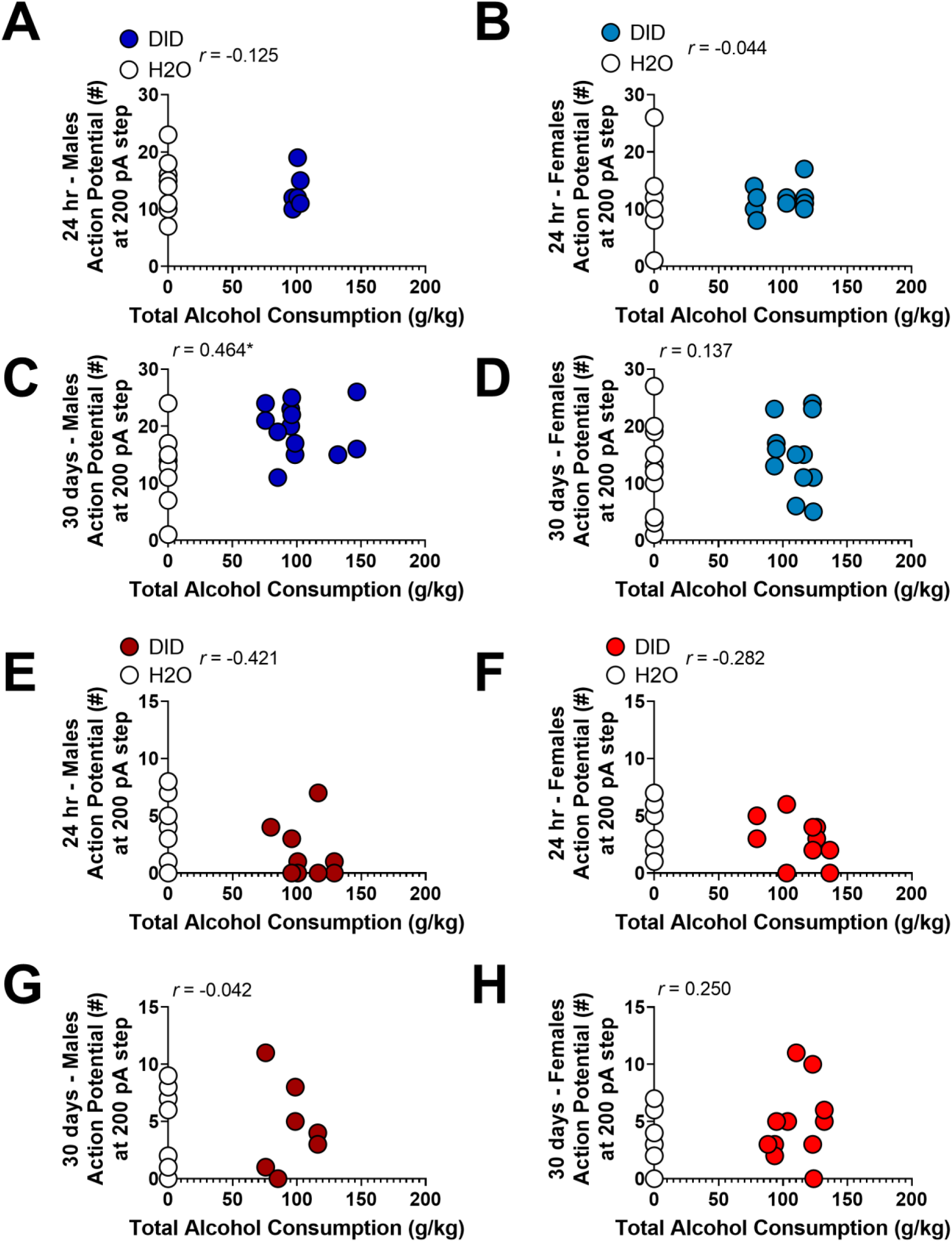
Total alcohol consumption during adolescent DID is correlated with SST hyperexcitability in adult male mice. (A-B) Total alcohol consumption across 4 cycles of adolescent DID was not significantly correlated with the number of action potentials fired during the final step of VI in SST neurons 24 hr after the end of DID. (C-D) Total alcohol consumption during adolescent DID is positively correlated with hyperexcitability in SST neurons in male mice, but not female mice, 30 days after the cessation of adolescent DID. (E-F) Adolescent alcohol consumption is not correlated with current-induced action potential firing in pyramidal neurons of male or female mice 24 hr after DID. (G-H) Total alcohol consumption during adolescent DID is not significantly correlated with pyramidal neuron action potential firing during the 200 pA step of VI in adult male or female mice. Filled circles represent individual cells from mice which underwent adolescent DID, with colors designating sex and cell type. Empty circles represent individual cells from control mice. * indicates *p* < 0.05.

## 4. DISCUSSION

Our work suggests that adolescent binge alcohol consumption via a modified DID model leads to modest but persistent alterations in cortical SST excitability (see **Figure 9** for overall model). We also provide preliminary evidence that this effect is not driven by HCN-channel alterations, as evident by no change in the action potential firing rate as measured by the ISI of SST neurons and no change in the voltage sag ratio in SST neurons. These general changes are consistent with a broad literature suggesting chronic adaptations in the cortex following adolescent models of alcohol exposure. Chronic ethanol administration induced sex-specific changes in the excitability of PL Martinotti neurons in adult rats, an effect mediated by differences in HCN channel function (Hughes et al., 2020). Changes in HCN channel function have also been reported in PL pyramidal neurons following adolescent alcohol exposure, leading to PL layer V pyramidal neuron hyperexcitability (Salling et al., 2018). However, we did not find changes in the ISI in SST neurons, suggesting that other mechanisms may also contribute to SST neuronal hyperexcitability. It is possible that HCN channel vulnerability is limited to deeper layers of the PL cortex. Future experiments should more explicitly explore HCN channel expression and function on SST neurons following adolescent alcohol.

**Figure 9.**
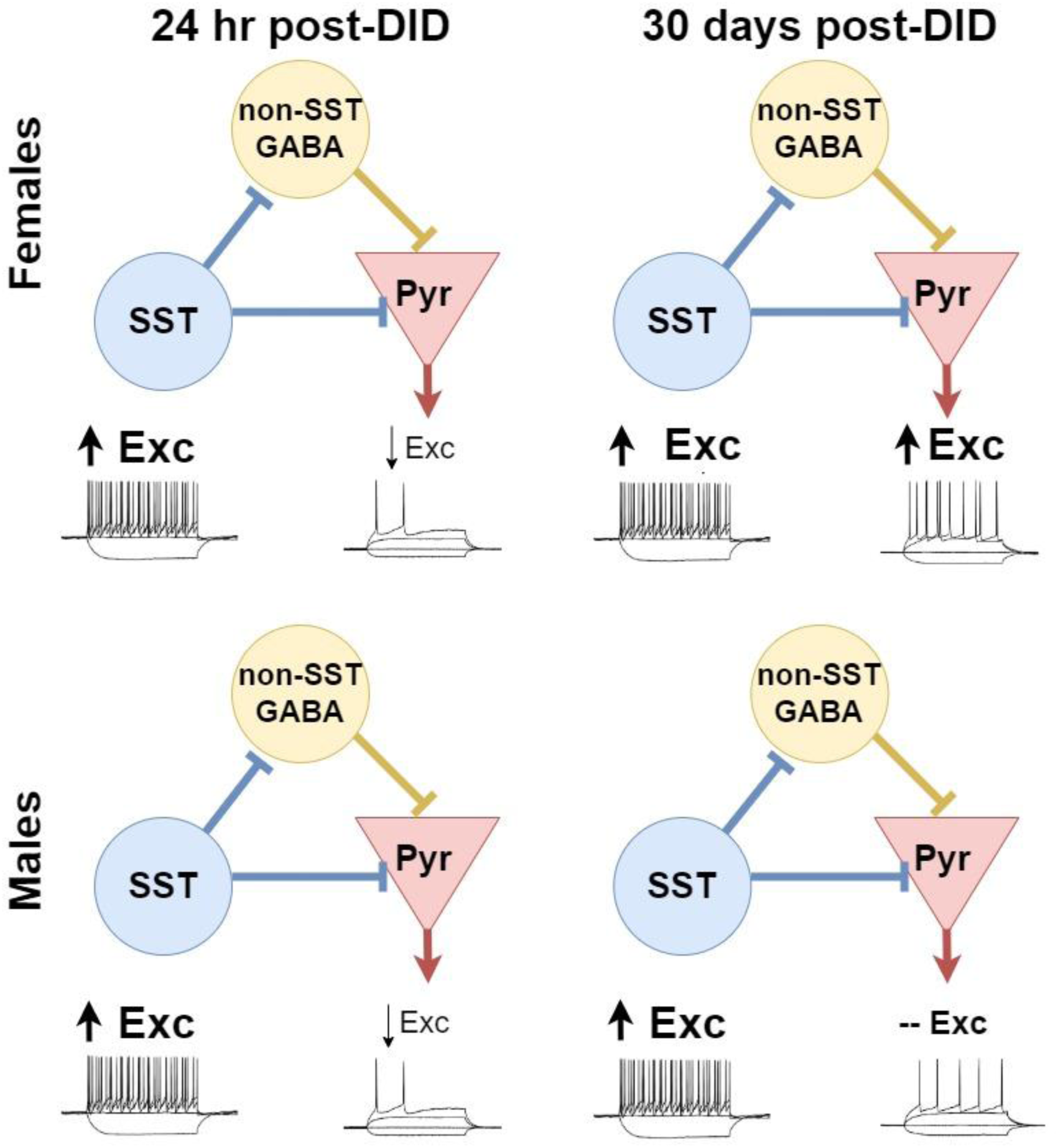
Overall model of microcircuitry changes 24 hr and 30 days after adolescent DID. Adolescent binge drinking disrupts the excitability of PL SST and pyramidal neurons, with changes in SST neurons persisting at least 30 days after cessation of drinking. In females, pyramidal neurons become hyperexcitable in adulthood. Despite these excitability changes, SST-mediated GABA transmission onto pyramidal and non-SST, non-pyramidal neurons is not altered 24 hr after ethanol consumption ends.

Importantly, differences in results seen at RMP and -70 mV could hint at voltage-gated channels as a key mechanism. Changes in L-type calcium channels are another potential mechanism for SST hyperexcitability following adolescent DID which we did not explore here. Treatment with ethanol increased the functionality of L-type calcium channels in cultured neocortical neurons (Katsura et al., 2006). L-type calcium channels modulate increased action potential firing in central amygdala neurons of non-ethanol-dependent rats (Varodayan et al., 2017). These channels have recently come into focus as a potential opportunity for pharmacological treatment of alcohol dependence (Little, 2021). So far, studies assessing L-type calcium channels after developmental alcohol exposure are limited.

As noted above, some studies have found persistent alterations in PL layer V pyramidal neuronal excitability after adolescent alcohol (Salling et al., 2018). In addition, intermittent exposure to alcohol during adolescence prevented the typical development of a tonic GABAergic current onto PL pyramidal neurons only in layer V (Centanni et al., 2017). Hyperexcitability may be most prominent in V pyramidal neurons, as others have shown that adolescent alcohol does not alter the intrinsic excitability of layer II pyramidal neurons in the short- or long-term in rats (Galaj et al., 2020). In our current study, we targeted layer II/III neurons in the PL cortex and found general increases in excitability weeks after the cessation of alcohol exposure. Galaj and colleagues used both a different species (rats) and a differing model of ethanol exposure (chronic intermittent ethanol exposure), highlighting the need for precise comparisons across differing models of alcohol exposure, and that models may not generalize across each other in their long-term effects (Crowley et al., 2019). Together, these findings all suggest that the consequences of alcohol exposure are age-dependent and may be layer-specific in the PL cortex, and can persist well into adulthood. We have shown here that adolescent binge drinking causes persistent alterations in PL SST neuronal excitability. Although we see largely similar effects in males and females, it is possible that the effects are driven by different mechanisms. Interestingly, this alteration in pyramidal neuron excitability appears to be driven by intrinsic mechanisms similar to that of SST neurons, as evident by a lack-of effect on optogenetically-evoked paired-pulse ratio from SST neurons onto either pyramidal and non-pyramidal neurons 24 hr after DID. Importantly, our excitability changes at 24 hr are more modest than those at 30 days, a timepoint we did not pursue with optogenetic methods.

Corresponding changes in SST peptide release following binge drinking remain under investigation. Although limited work has assessed changes in SST peptide release or expression in substance use disorders, SST represents a potential therapeutic target for a variety of neuropsychiatric illnesses (Brockway and Crowley, 2020). Future experiments should investigate changes in SST peptide release throughout development and following alcohol exposure using both optogenetic techniques (Dao et al., 2019), *in vivo* recording (Brockway et al., 2022, preprint), and novel biosensors (Wang et al., 2022, preprint; Xiong et al., 2021, preprint).

PL SST neurons have been implicated in anxiety-like behavior (Soumier and Sibille, 2014). Studies assessing changes in anxiety-like behavior following adolescent alcohol have found mixed results, with some showing increases in anxiety-like behavior after adolescent drinking (Lee et al., 2017; Varlinskaya et al., 2020) and others showing no change (Amodeo et al., 2018). While the current study did not explore the behavioral ramifications of long-term SST excitability, the literature has long suggested a broad role for these neurons in fear learning (Cummings and Clem, 2020), threat response (Joffe et al., 2022), and alcohol use itself (Dao et al., 2021), and we have recently shown their active engagement during exploratory behaviors (Brockway et al., 2022, preprint). Therefore, the consequences of a lifelong increase in SST function may have significant behavioral and neurobiological consequences.

There are several methodological and experimental design variables that contribute to the effects we uncovered. We used Ai9 reporter fluorescence, and expression of Ai32, as our primary marker of SST cells rather than strict electrophysiological properties such as high rheobase, hyperpolarized RMP, or low membrane resistance, as some studies have done (Joffe et al., 2020), as off-target expression of Cre recombinase in fast-spiking parvalbumin-expressing cells of SST-Cre mice has been reported previously (Hu et al., 2013). The SST-Cre line labels multiple SST subtypes, including non-Martinotti cells (Nigro et al., 2018), which we may have recorded from in this study. Although we did not use strict criteria values to exclude SST cells, recorded fluorescent cells which had a high membrane capacitance relative to membrane resistance or which fired wide action potentials (i.e. exhibited the previously described criteria for pyramidal neurons) were excluded. Our recording parameters are very similar to those reported in Joffe et al. 2020, highlighting consistency across groups.

We have previously reported, using identical DID and electrophysiology procedures, that binge drinking in adult mice reduces the excitability of PL SST neurons (Dao et al., 2021), in contrast to the present findings following adolescent binge drinking. This somewhat paradoxical effect is likely attributable to alcohol-induced disruptions in cortical development that are unique to alcohol exposure during adolescence. Importantly, little is known about the developmental trajectory of SST neurons – particularly within subregions of the prefrontal cortex. SST neurons are immature at the onset of adolescent DID (PND 28) and exposure to alcohol at this age alters the typical development of SST neurons. Previous work has demonstrated that the intrinsic excitability properties of SST neurons within other cortical regions (secondary motor cortex) continue to develop at least past PND 30 – and continue to influence monosynaptic connections between other neurons within the region as well (Pan et al., 2019). Others have shown similar development of SST neurons within somatosensory cortex (Kinnischtzke et al., 2012). This raises the likelihood that adolescent alcohol is disrupting the natural development of this pathway and leading to potentially long-term hyperexcitability, in contrast to the effects seen during adulthood. Therefore, our differences in adolescent versus adulthood (Dao et al., 2021) effects on SST neurons provide exciting insight into why adolescent drug consumption may be particularly problematic and uniquely alter the developing brain. In the infralimbic region of the prefrontal cortex, SST neuron input onto pyramidal neurons peaks in early adolescence and declines by early adulthood (Koppensteiner et al., 2019), possibly due to increasing SST cell density during early adolescence (Du et al., 2018). Synaptic pruning is another defining characteristic of rodent adolescent PFC development (Drzewiecki et al., 2016) that may be altered by adolescent binge drinking. Based on our results, we hypothesize that PL SST input onto pyramidal neurons increases and remains high following adolescent binge drinking, possibly as a result of impaired synaptic pruning. Interestingly, we do not see evidence of increased synaptic strength of SST projections onto pyramidal neurons (as measured by optogenetically-evoked PPR) suggesting that the elevated excitability of pyramidal neurons is not driven by increased SST drive onto them, but likely other intrinsic mechanisms. Others have proposed that adolescent alcohol exposure can interfere with proper PFC maturation processes, including those of GABAergic neurons (Centanni et al., 2017; Fish and Joffe, 2022). Much of the existing literature exploring the development of SST neurons focuses on either juvenile stages or adolescence separately, and studies characterizing SST neuronal properties throughout this early life window are limited. Additional studies investigating the typical development of SST neurons in the PL cortex are needed in order to fully understand how adolescent alcohol is interrupting this development.

## 4. CONCLUSIONS

Here, we provide increased validation for adolescent drinking in the dark as a clinically relevant model of binge ethanol consumption. We demonstrate that binge drinking during a broad definition of adolescence leads to increased excitability of PL SST neurons, which persists well into adulthood.

---

**Supplementary Figure 1.**
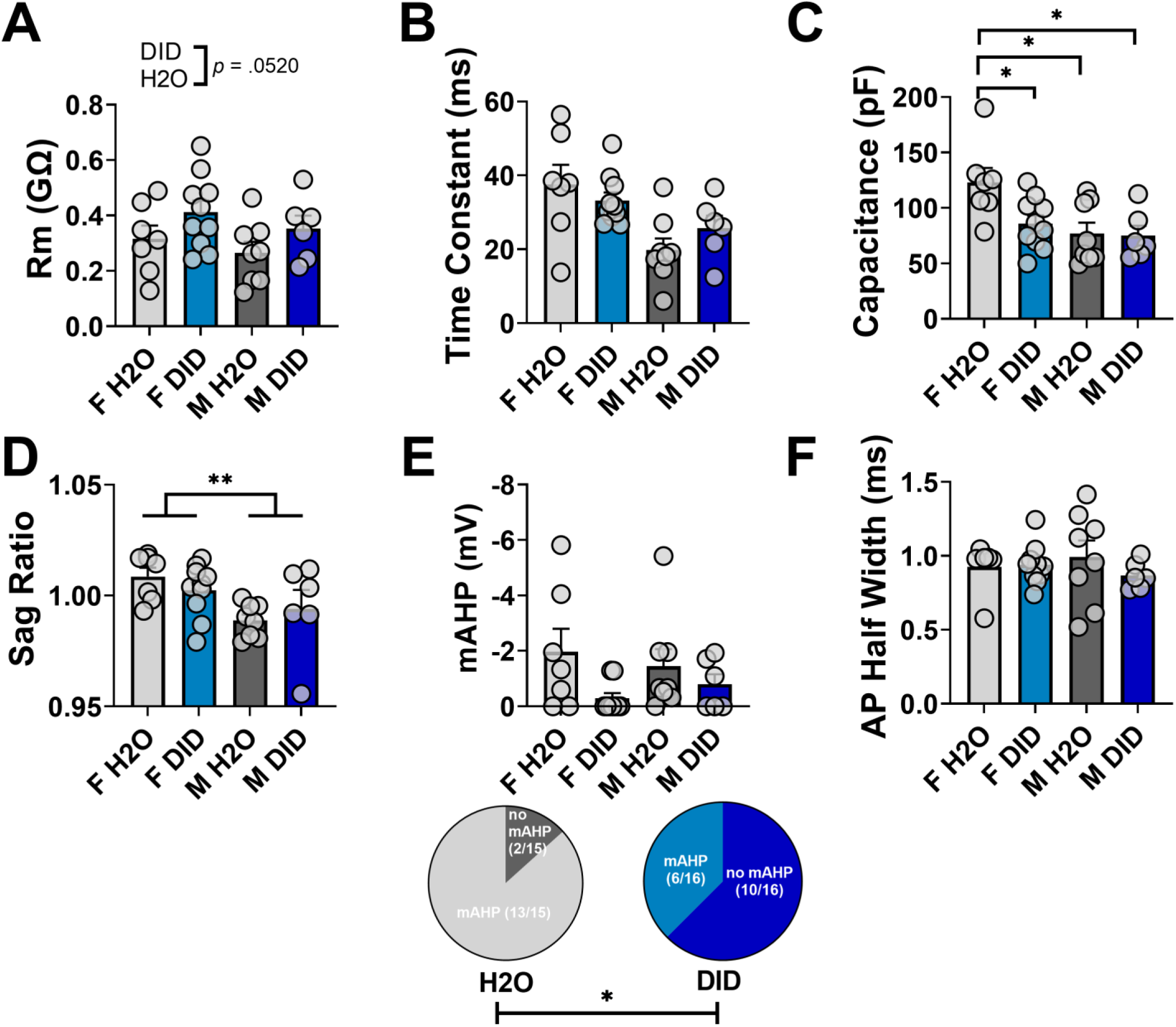
SST membrane properties 24 hr after adolescent DID. (A-B) There was no significant effect of adolescent DID or sex on input resistance or membrane time constant. (C) SST neurons from female H2O mice had a significantly higher capacitance than female DID, male H2O, and male DID mice. (D) SST neurons from female mice had a higher voltage sag ratio than SST neurons from male mice. (E) There was no effect of adolescent DID or sex on the mAHP of SST neurons. However, fewer neurons from DID mice exhibited mAHP following a 200pA injection step. (F) There was no effect of adolescent DID or sex on action potential half-width, measured in the first spike of rheobase.

**Supplementary Figure 2.**
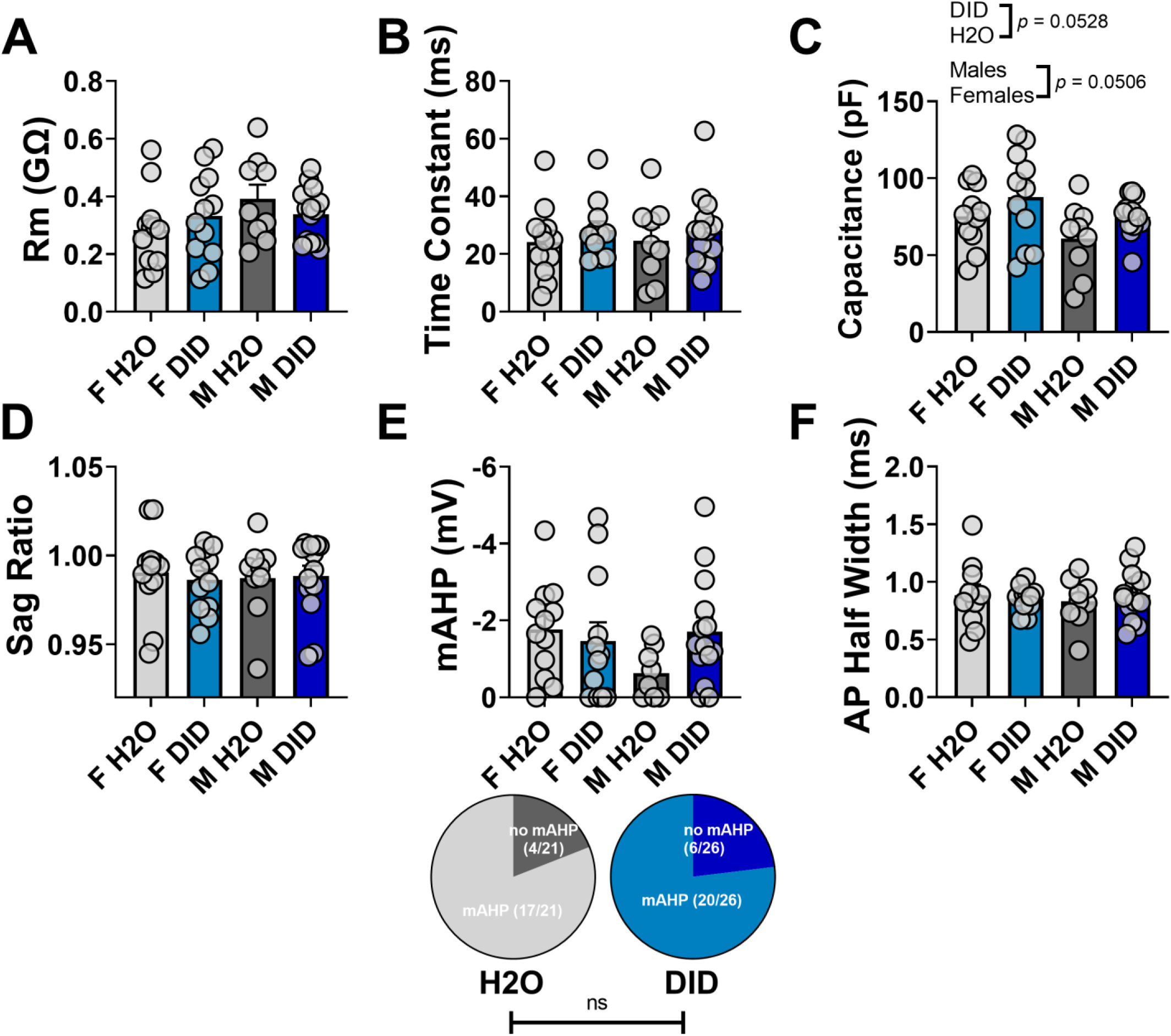
Membrane properties of SST neurons 30 days following the cessation of adolescent DID. (A-F) There was no effect of adolescent DID condition or sex on the input resistance, membrane time constant, membrane capacitance, voltage sag ratio, mAHP, or action potential half-width. Rm: membrane input resistance; mAHP: medium afterhyperpolarization

**Supplementary Figure 3.**
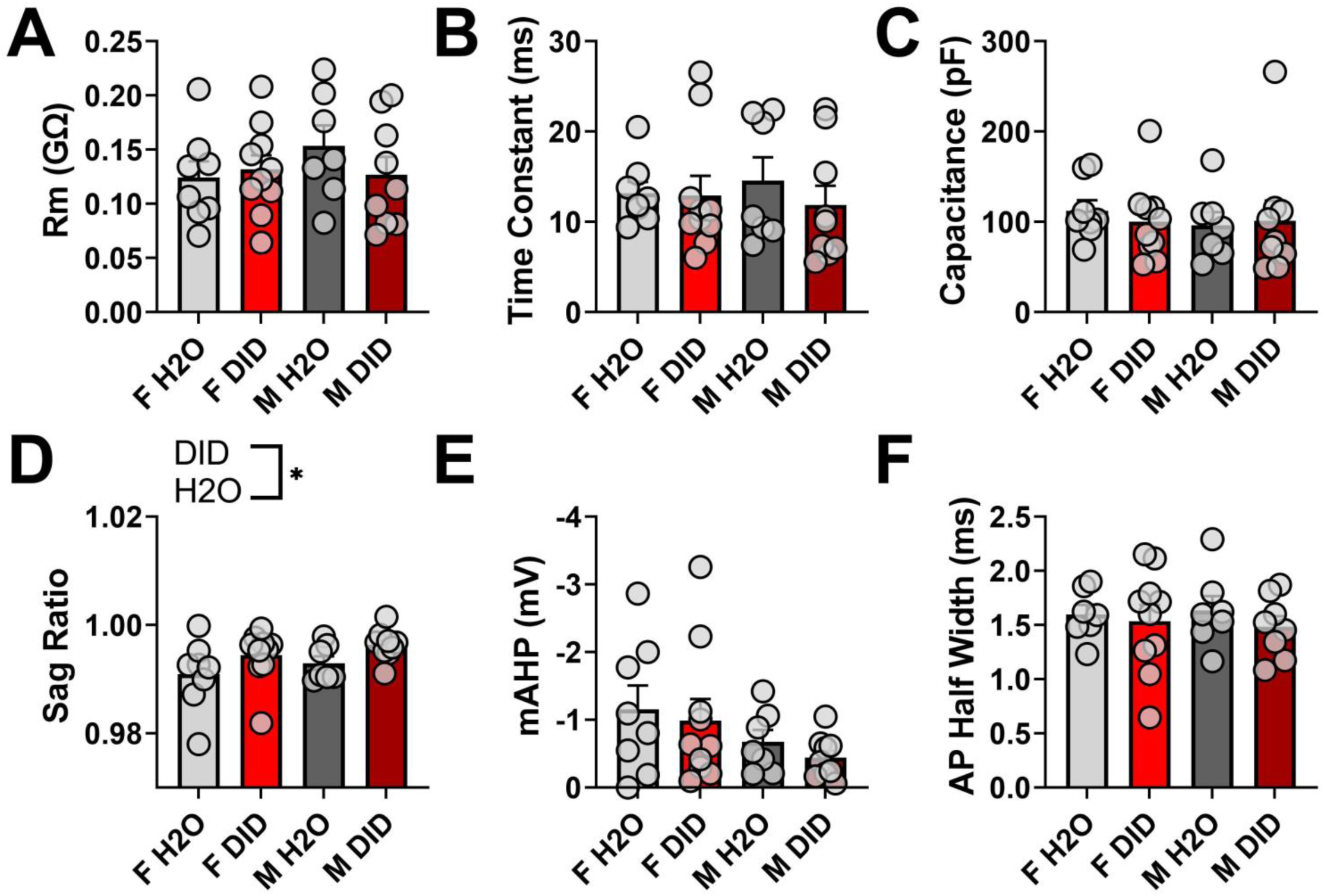
Membrane properties of pyramidal neurons 24 hr following the end of adolescent DID. (A-C) There was no effect of adolescent DID condition or of sex on input resistance, membrane time constant, or membrane capacitance. (D) There was a significant effect of adolescent DID on sag ratio. Sag ratios from pyramidal neurons from H2O mice were significantly lower than those of pyramidal neurons from DID mice, indicating that pyramidal neurons from H2O mice showed significantly more sag. (E-F) There was no change in mAHP or action potential half-width as a function of adolescent DID or of sex. Rm: membrane input resistance, mAHP: medium afterhyperpolarization.

**Supplementary Figure 4.**
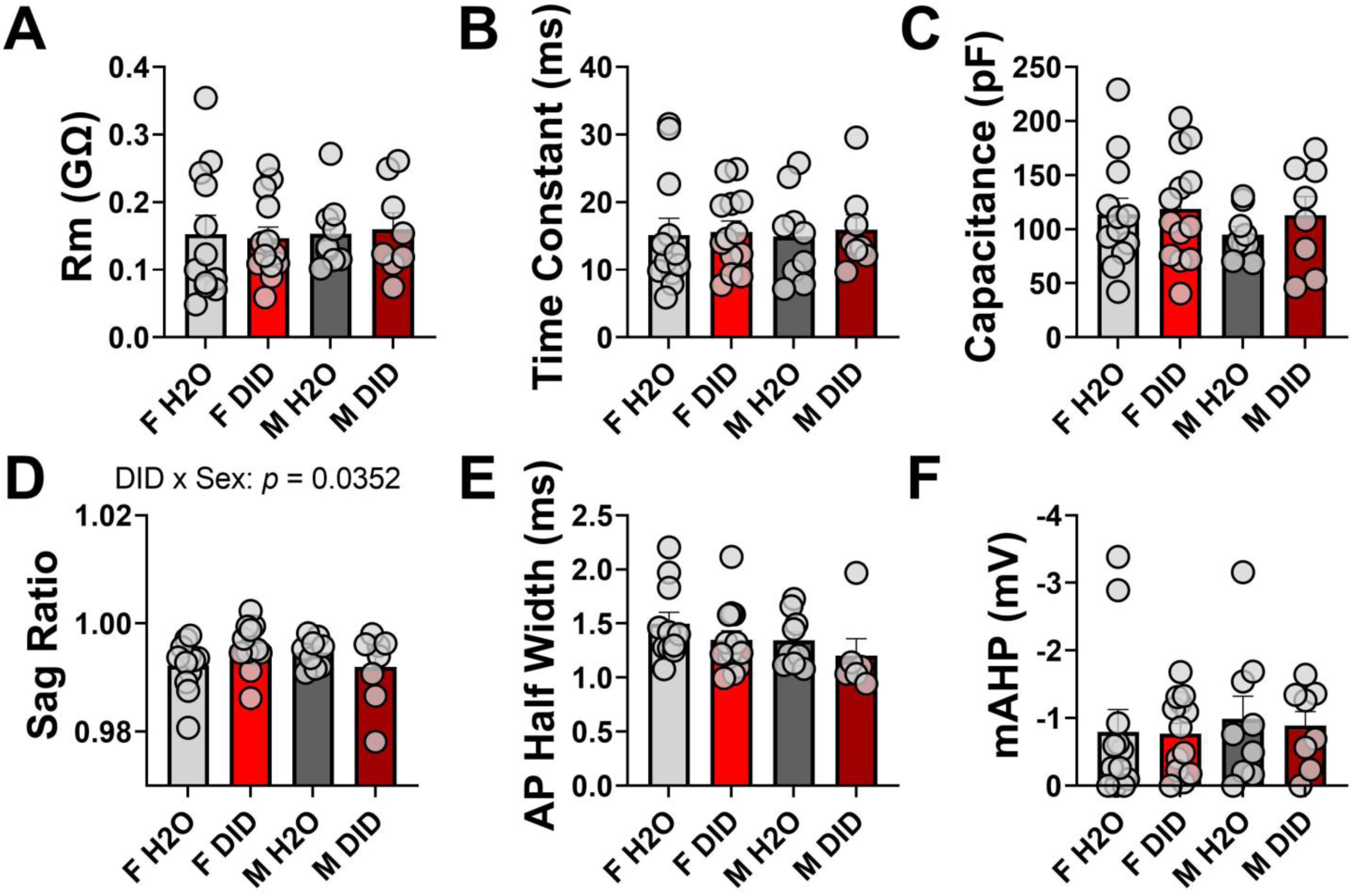
Membrane properties of PL pyramidal neurons 30 days after the end of adolescent DID. (A-C) There was no effect of adolescent DID condition or sex on input resistance, membrane time constant, or membrane capacitance. (D) In the voltage sag ratio, there was a significant DID x sex interaction. (E-F) Neither action potential half-width nor mAHP were altered as a function of drinking or sex in pyramidal neurons 30 days after adolescent DID concluded. Rm: input resistance, mAHP: medium afterhyperpolarization.

